# The encoding of interoceptive-based predictions by the paraventricular nucleus of the thalamus D2R+ neurons

**DOI:** 10.1101/2025.03.10.642469

**Authors:** Briana Machen, Sierra N. Miller, Al Xin, Carine Lampert, Lauren Assaf, Julia Tucker, Sarah Herrell, Francisco Pereira, Gabriel Loewinger, Sofia Beas

## Abstract

Understanding how the brain integrates internal physiological states with external sensory cues to guide behavior is a fundamental question in neuroscience. This process relies on interoceptive predictions, which are internal models that anticipate changes in the body’s physiological state based on sensory inputs and prior experiences. Despite recent advances in identifying the neural substrates of interoceptive predictions, the precise neuronal circuits involved remain elusive. In our study, we demonstrate that Dopamine 2 Receptor (D2R+) expressing neurons in the paraventricular nucleus of the thalamus (PVT) play key roles in interoception and interoceptive predictions. Specifically, these neurons are engaged in behaviors leading to physiologically relevant outcomes, with their activity highly dependent on the interoceptive state of the mice and the expected outcome. Furthermore, we show that chronic inhibition of PVT^D2R+^ neurons impairs the long-term performance of interoceptive-guided motivated behavior. Collectively, our findings provide insights into the role of PVT^D2R+^ neurons in learning and updating state-dependent predictions by integrating past experiences with current physiological conditions to optimize goal-directed behavior.

## INTRODUCTION

All organisms perceive the world through a combination of internal and external senses. As such, understanding how the brain integrates internal physiological states with external sensory cues to guide behavior is a fundamental question in neuroscience. Although much is known about perception of external sensory cues, less is known about the mechanisms mediating the perception of internal bodily cues. Interoception is the ability to sense, track, and integrate signals originating from within the body, such as hunger, thirst, temperature, heart rate, and blood CO_2_ levels, amongst others^1–5^. However, interoception plays a broader, more encompassing role within the systems governing emotion, motivation, decision making, and social cognition^4,6,7^. Moreover, impairments in the ability to monitor bodily signals exist across many psychiatric illnesses, such as panic disorder, anxiety, depression, and substance use disorders^2,3,8^.

Prevailing models of interoception posit that physiological signals are sensed, tracked, integrated, and prioritized to guide behaviors^1–6,8,9^. Studies on the mechanisms mediating interoception have largely focused on the “sensing” and “tracking” aspects. Indeed, hypothalamic areas, such as the arcuate nucleus (ARC) and the lateral hypothalamus (LH), as well as hindbrain areas, such as the medulla and nucleus of the solitary tract (NTS), play a distinctive role in detecting hunger signals from the body and the regulation of feeding behaviors^10,11^. Additional regions, such as the median preoptic nucleus (MnPO) and the subfornical organs (SFO), mediate thirst and water drinking behaviors^12–14^. However, much less is known about how the brain integrates these signals and prioritizes them to create the perception of internal states.

Once sensed, these peripheral physiological signals are integrated with external sensory cues and information from prior experiences to create internal models^4,5,9^. These internal models enable individuals to anticipate changes in the body’s physiological state based on these factors (current physiological state, environmental cues, learned experiences) – a process known as interoceptive prediction^4,5,9^. These predictive processes allow the brain to prepare for and respond proactively to physiological demands, ensuring efficient and adaptive regulation of bodily processes. Such mechanisms also represent a fundamental aspect of how the brain maintains homeostasis and guides behavior. By predicting physiological needs, the brain prioritizes actions that restore or sustain internal balance, ultimately shaping decision-making and motivation. While significant advances have been made in identifying the neural substrates underlying interoceptive predictions, the specific neuronal circuits responsible for encoding these processes remain poorly understood.

The paraventricular nucleus of the thalamus (PVT) has emerged as a critical node in the limbic system, yet its functional role remains unclear. Anatomical studies show that the PVT receives converging inputs from brain areas that process homeostatic and interoceptive signals^15–17^. Hypothalamic and brainstem areas that process physiological signals, such as hunger^11,18–20^, thirst^12–14^, thermoregulation^21,22^, pain^23–25^, and stress^26–28^ project to and recruit the PVT. Moreover, optogenetic activation of inputs to the PVT from hunger processing areas, such as the ARC and LH, result in feeding and food-seeking behaviors^18,20,29^; while stimulation from inputs originating in the MnPO result in water intake and seeking behavior^13,14^. These studies demonstrate that PVT neurons participate in interoception. In addition, the PVT receives signals about environmental cues from cortical areas, such as the medial prefrontal cortex (mPFC)^30–34^. The activity of PVT neurons is also altered by external cues when paired with physiologically relevant stimuli^18,27,30,32–36^. As such, the PVT is anatomically well-positioned to effectively bridge the interoceptive and exteroceptive domains to create internal states that guide behavior.

While commonly glutamatergic, PVT projection neurons are heterogeneous^26,37–39^. Particularly, our previous research found that a subtype of PVT neurons (PVT neurons expressing the dopamine receptor 2; PVT^D2R+^) are tuned to changes in current physiological states^26,39^ and, as such, play a more general role in integrating and encoding these states. In this study, we used fiber photometry to further investigate the *in vivo* dynamics of PVT^D2R+^ neurons in mice performing a reward linear maze task. Specifically, we focused on thoroughly characterizing the precise role these neurons play in interoceptive perception that influence goal-directed behaviors. We discovered that PVT^D2R+^ neurons integrate factors, such as physiological needs, environmental cues, and previous experiences, to compute an interoceptive prediction. This prediction, in turn, guides motivated behavior. Moreover, we show that chronic chemogenetic inhibition of PVT^D2R+^ neurons results in long-term deficits in interoceptive guided motivated behavior. Collectively, our results highlight the reconceptualization of PVT^D2R+^ neurons as a neural mechanism that is critical for interoception and interoceptive predictions that are needed for the perception of internal states.

## METHODS

### Mice

All procedures were performed in accordance with the University of Alabama Birmingham *Institutional Animal Care and Use Committee (IACUC)* Guide for the Care and Use of Laboratory Animals. For this, female and male Drd2-Cre (GENSAT founder line ER44) or C57BL/6 mice were used. These mice were between 8 – 15 weeks of age and were group housed under a 12-h light-dark cycle (6 a.m. to 6 p.m. light), at a temperature of 70–74 °F and 40–65% humidity. After surgery, mice were single-housed and were provided with food and water *ad-libitum.* Two days before initiating behavioral training procedures, food was removed for food restriction and the amount of food provided was according to 85% of their free-feeding weight. Water was maintained *ad-libitum,* unless otherwise specified. Mice returned to *ad libitum* feeding after testing.

### Viral vectors

For photometry experiments, we used AAV9-hSyn-FLEX-GCaMP6s-WPRE-SV40 which was produced by the Vector Core of the University of Pennsylvania. For our chemogenetic inhibition, we used AAV2-hSyn-DIO-hM4D(Gi)-mCherry which was produced by Addgene. For controls, we used AAV2-CAG-Flex-tdTomato which was produced by the vector core of the University of North Carolina. GABA sensors were AAV-Syn-iGABASnFR2 or AAV-Syn-Flex-iGABASnFR2^40^ purchased from Addgene.

### Stereotaxic surgery

Mice were given viral injections (approximately 1 μl) at the following stereotaxic coordinates (at a 6° angle): pPVT, −1.60 mm from Bregma, 0.25 mm lateral from midline, and −3.35 mm vertical from the cortical surface. Optical fibers with diameters of 400 μm (0.48 NA - Neurophotometrics) were used. These fibers were implanted over the pPVT immediately following viral injections (targeted 300-400 μm above the injection site). Following stereotaxic injections, animals were allowed to recover for 2–3 weeks to ensure maximal AAV expression.

### Linear maze reward-seeking task

The linear maze task consisted of a linear maze (81.28 × 21.59 × 29.21, L ×W× H in cm). The Ethovision (Noldus) tracking system allowed us to track the movement of the mice through the maze. The maze consisted of three zones: trigger zone, corridor, and reward zone. The opposite ends of the track were designated as the trigger zone and reward zone and were connected by a long corridor. The task consisted of 60 min-long sessions of self-paced trials and mice received one test session per day. For each trial, food-restricted mice were trained to wait in the trigger zone for two seconds. An LED mounted on top of the reward magazine port was then illuminated, which signaled reward availability, allowing mice to run from the trigger zone down the corridor and into the reward zone to retrieve a food reward. The strawberry milkshake (SMS) reward consisted of strawberry Ensure® mixed 1:1 with 20% sucrose water. The non-caloric reward consisted of 10% sucralose (Splenda®) solution. The sucrose reward consisted of 10% sucrose solution. Reward delivery and food consumption signaled trial termination; therefore, mice had to return to the trigger zone to initiate another trial. Experimental schedule and data acquisition were implemented through the Ethovision software for operant control. For training in this task and to avoid potential neophobia, mice were provided with 8 hours of SMS reward in their home cages for two consecutive days before training began. For the initial training session, mice performed the linear maze task without a wait requirement in the trigger zone, so no premature trials occurred. In subsequent sessions, a rule was implemented requiring mice to remain in the trigger zone for two seconds before approaching, allowing for baseline activity measurement prior to approach.

### Blocked reward access sessions

Mice that had previously been trained on the linear maze reward-seeking task underwent testing in a modified session in which access to the reward port was physically obstructed. Each session comprised two distinct phases: a 15-minute *Block phase* followed by a 60-minute *Post-Block phase*. During the *Block phase*, a wire mesh tea strainer was used to block access to the reward port. Upon completion of this phase, the session was briefly paused to remove the wire mesh, after which the *Post-Block phase* commenced and continued uninterrupted for the remaining duration. These sessions were conducted as self-paced trials, with no rewards delivered at any point during the session.

### Reward magnitude test sessions

Mice previously trained on the linear maze reward-seeking task were tested in a modified session where the amount of reward varied between sessions. Each 60-minute session consisted of three consecutive blocks of 33 trials, with each block delivering a different reward amount: a small reward (1 µl), a medium reward (3 µl), and a large reward (6 µl). Mice were required to complete the small reward trial block before proceeding to the medium reward block, followed by the large reward block.

### Bulk Ca^2+^ and fiber photometry

All photometry experiments were performed using an RZ5P acquisition system (Tucker-Davis Technologies; TDT) equipped with a real-time signal processor and controlled by a software interface (Synapse version 92). Specifically, the system is integrated with two continuous sinusoidally modulated LEDs at 473 nm (211 Hz) and 405 nm (531 Hz) that serve as the light source to excite GCaMP6s and an isosbestic autofluorescence signal, respectively. The LED intensity (ranging 10–15 μW) at the interface between the fiber tip and the animal was constant throughout the session. Fluorescence signals were collected by the same fiber implant that was coupled to a 400 μm optical patch-cord (0.48 NA) and focused on two separate photoreceivers (2151, Newport Corporation) connected to the emission ports of a custom-ordered fiber photometry Mini Cube (Doric Lenses). TTL pulses recorded by the same system were used to annotate the occurrence of behavioral manipulations. For the measurements of fluorescent calcium signals and ΔF/F analysis, a least-squares linear fit to the 405 nm signal to align it to the 470 nm signal was first applied. The resulting fitted 405 nm signal was then used to normalize the 473 nm as follows: ΔF/F = (473 nm signal − fitted 405 nm signal)/fitted 405 nm signal. All GCaMP signal data is presented as the z-score of the ΔF/F from a baseline (2 seconds) prior to the onset of events.

### Four-bottle choice assessment

Mice were individually housed and tested in their home cages. After becoming accustomed to each solution, hungry mice were given free access to four bottles in their home cage. Each bottle contained one of the following: water, a 10% sucrose solution, a non-caloric 10% sucralose solution, or a strawberry milkshake (strawberry Ensure® mixed with 20% sucrose water). After a 2-hour testing period, the total volume consumed (in mL) for each solution was recorded. Consumption was averaged across two sessions. To prevent place preference, the placement of the solutions in the cage was randomized across sessions.

### Chemogenetic inhibition

For chemogenetic inhibition, deschloroclozapine (DCZ; Tocris Bioscience, catalog #7193) was dissolved in 1–2% dimethyl sulfoxide (DMSO) in saline and given intraperitoneally (I.P.) at a final dose of 0.1 mg/kg. An equivalent volume of 1-2% DMSO saline was used as vehicle control on SMS pre-exposure days and on post-inhibition testing days. Alternatively, CNO (Enzo Life Sciences, catalog# BML-NS105) was administered at a dose of 10 mg/kg. All injections were administered 30 minutes prior to behavioral testing.

### *In vivo* data analysis

After calculating the z-score of the ΔF/F for each event in every trial the mice performed, the latency to the reward zone was calculated. All trials that took longer than 11 seconds and the events corresponding to that trial (decision to approach, approach, reward zone arrival, reward delivery) were excluded. Less than 10% of the trials met this exclusion criterion. After that, the trials and events for each individual test session were sorted by their trial order within a session. After sorting the trials in each test session, these were then divided into 5 equivalent trial blocks. This binning strategy allowed for comparable representation per animal in both block trials and was used by prior studies to test the effect of trial order^39^, which permits the comparison of the activity dynamics of PVT neurons and how they were modulated by the motivational states of the mice across the progression of each test session.

*FLMM* - Trial order effects were also confirmed using a Functional Linear Mixed Model (FLMM). This analysis, previously described in detail^39,41,42^, combines linear mixed models (LMM) and functional regression. The FLMM framework produces plots for each covariate in the model (e.g., trial order effects) to show whether associations with the photometry signal are significant at each timepoint during the behavior. These FLMM output plots display the regression coefficient estimates for each covariate at each timepoint in the trial (solid lines). They also show estimated pointwise (dark-shaded area) and joint (light-shaded area) 95% confidence intervals (CI) for the coefficient estimates. The joint CIs allow for the interpretation of each covariate’s effect on the photometry signals across multiple time intervals in the trial. Timepoints where the pointwise CI do not include zero indicate pointwise statistical significance, while timepoints where the joint CI do not include zero indicate joint statistical significance. For the trial order effects we used the FLMM association with continuous variables (https://rpubs.com/gloewinger/1159129). The random intercept model: IE[*Y_i,j_*(*s*)| *X_i,j_*, γ*_i_(s)] =* β*_0_(s) + X_i,j_* β*_1_(s) +* γ*_0,i_(s)* was chosen based on best fit determined by their AIC (Akaike Information Criterion) and BIC (Bayesian Information Criterion) values.

*Concurrent FLMM Model* – We applied a concurrent FLMM model to analyze trials of different lengths and estimate how factors that vary within a trial (e.g., state) influence the photometric signals of PVT^D2R+^ neurons. This was necessary as animals took different amounts of time to travel between the trigger and reward zones and thus each “trial” could differ in length. Specifically, we included an “approach” variable (a binary indicator of whether an animal was approaching the reward zone at each trial timepoint) in the model, in addition to its interaction with the trial outcome type. For animal *i* on trial *j,* denote:

- *X_1,I,j_:* the trial outcome, where *X_1,I,j_* = 1 for ‘strawberry milkshake’ (SMS) trials and *X_1,I,j_* = 0 for ‘other outcome’ trials.
- *X_2,I,j_(s)*: approach, where *X_2,I,j_(s)* = 1 when mouse *i* is approaching the reward zone on trial *j* at trial timepoint *s* and *X_2,I,j_*(s)= 0 for trial timepoints after the reward has been delivered.
- *Y_i,j_(s)* is the photometry response of mouse *i* on trial *j* at trial timepoint *s*.

β*(s)* is the vector of coefficients at timepoint *s* and L*_i,j_(s)* is Gaussian noise for mouse *i* on trial *j* at trial timepoint *s*.

We used the formula syntax: photometry ∼ outcome * approach + (outcome |ID). The model formula was:

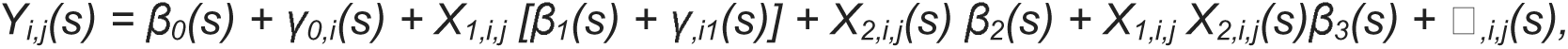

where (*s*) indicates that a variable is functional (i.e., varies across trial timepoints). *X_1,i,j_* does not vary with respect to *s*, as it denotes a trial condition.

The coefficients are interpreted:

- β*_0_(s)*: The mean signal when a mouse has received (i.e., after reward delivery) an “other” outcome (e.g. a non-caloric reward) at trial timepoint *s*.
- β*_1_(s)*: The mean difference in signal between hungry strawberry milkshake reward (H-SMS) and the other outcome (e.g. a non-caloric reward) during reward approach (i.e., before non-caloric reward delivery) at trial timepoint *s*.
- β*_2_(s)*: The mean difference between approach and outcome delivery for other outcome trials at trial timepoint *s*.
- β*_3_(s)*: This interaction term is interpreted as a difference in differences. Denoting *A(s)* as the mean signal difference between approach and reward in H-SMS (at trial timepoint *s*) trials, and *B(s)* as the mean signal difference (at trial timepoint *s*) between approach and reward in the outcome of comparison (e.g. a non-caloric reward), β*_3_(s)* = *A* − *B*. Thus, β*_3_(s)* = 0 when the magnitudes of the approach vs. reward zone mean signal difference are equal (at timepoint *s*) on H-SMS and “other” outcome (e.g. a non-caloric reward) trials. Conversely, β*_3_(s)*≠ 0 when approaching and H-SMS are together associated with an additional effect. Thus, β*_3_(s)* captures the degree to which the trial outcome type (e.g. H-SMS vs. non-caloric rewards) modulates the change in signal that occurs when an animal receives the reward (compared to approaching the reward zone).

Altogether, this concurrent FLMM model enables a detailed understanding of how context (state and outcome) and behavioral phase (approach vs. outcome delivery) dynamically influence neuronal activity, shedding light on the differences in anticipatory signaling during motivated behavior.

*Linear Interpolation-* Linear interpolation was used to estimate unknown values that fall within the range of known data points using the following mathematical formula:

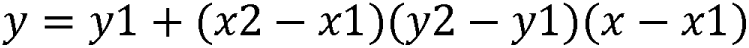

In this study, linear interpolation was employed to standardize the length of neuronal recording data across trials of varying durations. This method estimates intermediate values between known data points by assuming a linear relationship. Specifically, here, the recording of each trial was resampled to the same number of data points using linear interpolation, preserving the overall temporal structure while mitigating the effects of varying trial durations. This strategy allowed for consistent alignment and aggregation of neuronal activity patterns across trials. Linear interpolation was solely used to generate average traces for graphical purposes showing photometry signals during approach (Fig. 4A-F). These approach signals were analyzed by calculating the AUC of the signal during approach timepoints: for trial *j,* we summed the signal values during approach timepoints and divided that sum by the number of approach timepoints). We then conducted standard hypothesis tests on these AUCs.

### Histology

To verify GCaMP6s, GABASnFR2, hM4Di, and TdTomato expression and optical fiber placements, mice were injected with euthanasia solution and subsequently sacrificed via transcardial perfusion, first with PBS and then with paraformaldehyde (PFA; 4% in PBS). Brains were then post-fixed in 4% PFA at 4 °C overnight and cryoprotected using a 30% PBS-buffered sucrose solution for ∼24–36 h. Coronal brain sections (50 μm) were generated using a freezing microtome. Images were taken using a Nikon microscope running NIS Elements AR 5.42.03 64-bit (Nikon).

### Statistics and data presentation

All data were analyzed using GraphPad Prism (Domatics) and custom-made Python scripts (Anaconda Navigator). FLMM analyses were performed using R scripts (R Studio). For the FLMM Trial Order Effects, the associations with continuous variables and random intercept model was used^39,41,42^. For statistical analyses, two-tailed paired and unpaired t-tests (parametric; test statistic: t), repeated measures ANOVAs, and two-way ANOVAs (parametric; test statistic: F) were used. Post-hoc Tukey’s, Dunnett’s, and Šídák’s multiple comparisons test were used when appropriate and if the corresponding omnibus test detected a significant difference. All data are presented as mean ± s.e.m.

## RESULTS

### PVT^D2R+^ neurons sense and track hunger states

We first examined how physiological states impacted PVT^D2R+^ neuronal activity while mice performed a linear maze task that assesses motivation. For this, we trained hungry mice to perform a linear maze task while investigating the *in vivo* dynamics of PVT^D2R+^ neurons (Fig. 1A,B). In this task, mice were placed in a linear maze equipped with a reward delivery port and a bright cue LED light at the end wall of the maze (reward zone). The other far end of the maze was set as the “trigger zone” (Supp. Fig. 1A). Mice had to learn by trial and error to perform trials requiring them to go and remain in the trigger zone for 2 sec to initiate a rewarded trial, which was signaled by a cue light above the reward port. Once the cue light signaled reward availability, mice could approach the reward zone to obtain a reward (strawberry milkshake, SMS). We found that when mice approached the reward zone, PVT^D2R+^ neuronal activity significantly increased (Supp. Fig. 1B). In accordance with their previously suggested role in sensing interoceptive stimuli, such as hunger, we found that the hunger states modulated PVT^D2R+^ neuronal activity. To show this, we first sorted trials performed by trial order, labeling the earliest trials of the session as ‘Trial Group - G1’ and the latest trials as ‘G5’^39^ (Supp. Fig. 1C). We implemented this strategy to model the assumption that well-trained food-restricted mice are hungrier at the beginning of each 60 min test session than at the end (when they have eaten several rewards). Importantly, here, the average latency to approach did not differ across the trial groups throughout test sessions (Supp. Fig. 1D). When comparing between early (G1) and late (G5) trials, the mean PVT^D2R+^ neuronal responses were significantly higher in the early trials (Supp. Fig. 1E,F), suggesting that stronger hunger levels lead to higher PVT^D2R+^ neuronal activity. We also found a strong negative correlation between the area under the curve (AUC) during approach and trial order (Supp. Fig. 1G), such that higher PVT^D2R+^ neuronal activity was associated with early trials and diminished activity with later trials. The association between trial order and decreases in signal of PVT^D2R+^ neurons was confirmed using a functional linear mixed modeling (FLMM) statistical framework^39,41,42^ (Supp. Fig. 1H). Notably, no correlation was found between approach AUC and latency to approach (Supp. Fig. 1I).

**Figure 1.**
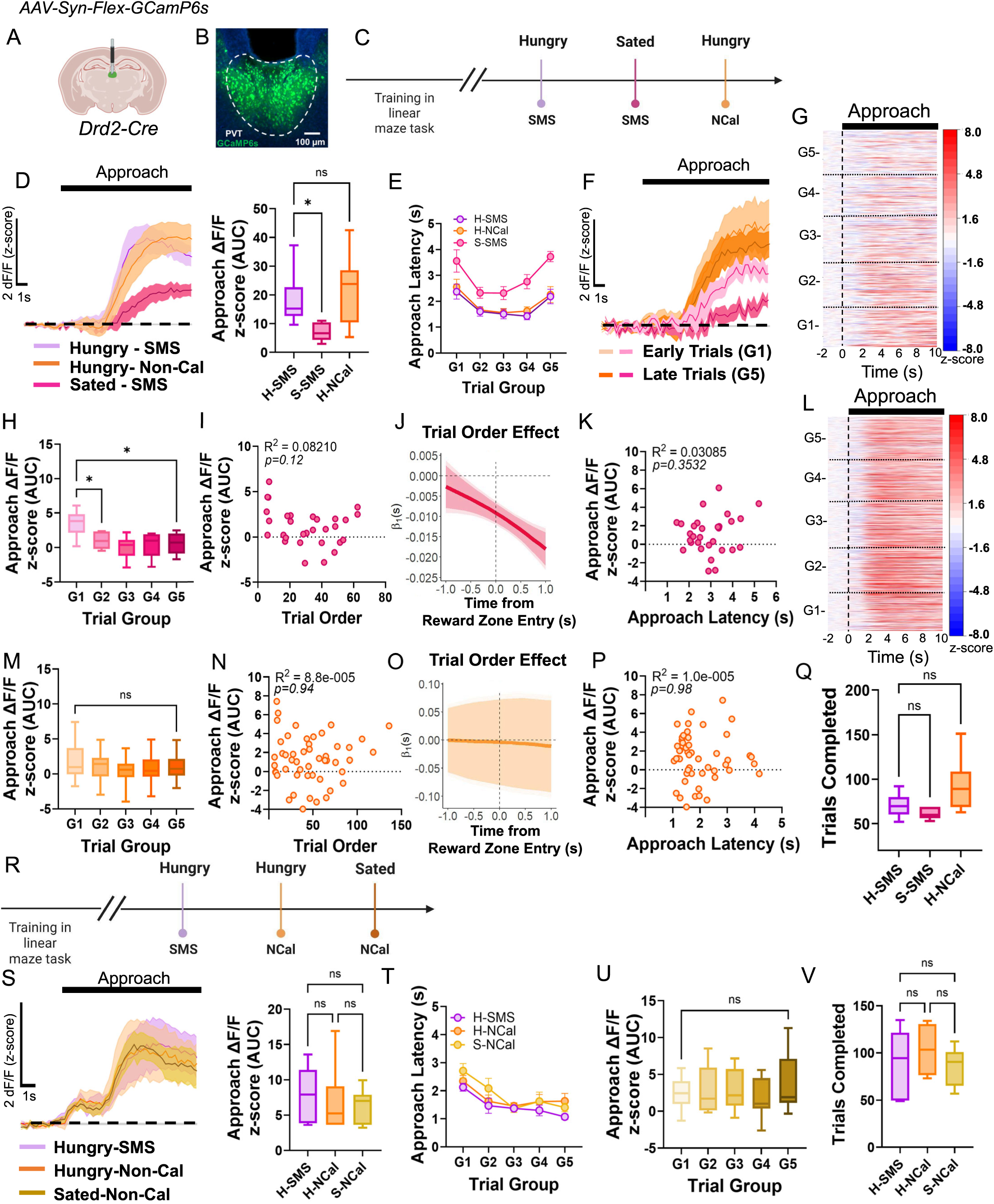
PVT^D2R+^ neuronal activity state-dependent modulation is contingent on calorie intake. **(A)** Schematic of the stereotaxic injections and optical fiber implantation for targeting D2R+ neurons in the PVT. **(B)** Representative image from a Drd2-Cre mouse expressing GCaMP6s in the PVT. **(C)** Timeline schematic for the test sessions. **(D)** Average GCaMP6s responses (n=5) from PVT^D2R+^ neurons and AUC quantification comparing ramping activity during reward approach in test sessions: H-SMS: n = 705 trials, S-SMS: n = 368 trials, and H-NCal: n = 928 trials. Repeated measures ANOVA, *p<0.05. H-SMS vs. S-SMS Dunnett’s multiple comparisons test *\*p<0.05*. H-SMS vs. H-NCal Dunnett’s multiple comparisons test *p=0.54, ns*. **(E)** Latencies to reach the reward zone across ‘trial group blocks’ (G1 – G5) for H-SMS and S-SMS sessions. Two-way ANOVA, Effect of trial group block *\*\*\*\*p<0.*0001; Effect of test session (satiety) *\*\*\*\*p<0.0001;* Interaction *p=0.46*, *ns*. H-SMS and H-NCal sessions. Two-way ANOVA, Effect of trial group block ****p*<*0.0001; Effect of test session *p=0.44*, *ns*; Interaction *p=0.99, ns*. **(F)** Average GCaMP6s responses from PVT^D2R+^ neurons comparing reward approach between early ‘G1’ and late ‘G5’ trials within the S-SMS and the H-NCal test sessions. **(G)** Heatmap of reward approach responses from PVT^D2R+^ neurons during sated-SMS test sessions, time-locked to approach onset, sorted by trial order, and binned into 5 ‘trial group blocks’ (G1 – G5). **(H)** AUC quantification of the reward approach-evoked changes in GCaMP6s activity across trial group blocks within the sated-SMS test sessions. Repeated measures ANOVA, *\*\*p<0.001*. G1 vs. G2 Dunnett’s multiple comparisons test *\*p<0.05*. G1 vs. G5 Dunnett’s multiple comparisons test *\*p<0.05*. **(I)** Correlation between the AUC of the reward approach-evoked changes in GCaMP6s activity and trial order within the sated-SMS test session. **(J)** FLMM coefficient estimates plot of the approach trial order effect and statistical significance at each trial time-point results for the photometric responses of PVT^D2R+^ neurons for reward zone entry. The plot shows a negative association between PVT^D2R+^ GCaMP6s responses and trial order before and after reward zone entry for the S-SMS test session. **(K)** No correlation between the AUC of the reward approach-evoked changes in GCaMP6s activity and approach latency within the S-SMS test session. **(L)** Heatmap of reward approach responses from PVT^D2R+^ neurons during H-NCal test sessions. **(M)** AUC quantification of the reward approach-evoked changes in GCaMP6s activity across trial group blocks within the H-NCal test sessions. Repeated measures ANOVA, *p=0.078, ns*. G1 vs. G5 Dunnett’s multiple comparisons test *p=0.43, ns*. **(N)** No correlation between the AUC of the reward approach-evoked changes in GCaMP6s activity and trial order within the H-NCal test sessions. **(O)** FLMM coefficient estimates plot of the approach trial order effect and statistical significance at each trial time-point results for the photometric responses of PVT^D2R+^ neurons for reward zone entry. The plot shows no significant association between PVT^D2R+^ GCaMP6s responses and trial order for the H-NCal test sessions. **(P)** No correlation between the AUC of the reward approach-evoked changes in GCaMP6s activity and approach latency within the H-NCal test sessions. **(Q)** Quantification of trials completed within a session comparing trials during the H-SMS, the S-SMS and the H-NCal test sessions. Repeated measures ANOVA *\*p<0.05.* H-SMS vs. S-SMS Dunnett’s multiple comparisons test *p=0.26, ns.* H-SMS vs. H-NCal Dunnett’s multiple comparisons test *p=0.06, ns.* **(R)** Schematic of the timeline for the test sessions for the second cohort of mice. **(S)** Average GCaMP6s responses (n=4) from PVT^D2R+^ neurons and AUC quantification comparing ramping activity during reward approach in test sessions: H-SMS, H-NCal, and S-NCal. H-SMS: n = 540 trials, H-NCal: n = 414 trials and S-NCal: n = 515 trials. Repeated measures ANOVA, *p=0.86, ns*. **(T)** Latencies to reach the reward zone across ‘trial group blocks’ (G1 – G5) for H-SMS, H-NCal, and S-NCal sessions. Two-way ANOVA, Effect of trial group block *\*\*\*\*p<0.*0001; Effect of test session \**p<0.05;* Interaction *p=0.82, ns*. **(U)** AUC quantification of the reward approach-evoked changes in GCaMP6s activity across trial group blocks within the S-NCal test sessions. Repeated measures ANOVA, *p=0.83, ns*. **(V)** Quantification of trials completed during H-SMS, H-NCal and S-NCal test sessions. Repeated measures ANOVA, *p=0.64, ns*.

To further probe the ability of PVT^D2R+^ neurons to sense and track hunger levels, we recorded the responses of these neurons during two separate additional test sessions (Fig. 1C) in which: 1) sated mice received a calorie-rich SMS reward (Sated-SMS; S-SMS), and 2) hungry mice received a sweet non-caloric reward (Hungry-Non-Cal). Importantly, we used a within-animal comparison strategy to circumvent potential confounds due to differences in viral expression or fiber placement. For these test sessions, we hypothesized that sated mice would display an overall decrease in PVT^D2R+^ neuronal activity levels across all trials in the session. Moreover, we also hypothesized that if PVT^D2R+^ neurons tracked the hunger levels of the mice when receiving a non-caloric reward (H-NCal), this would result in hungry mice remaining hungry, and thus, PVT^D2R+^ neuronal activity would not be modulated across the entire test session. Indeed, we observed that when mice were tested in a sated state, there was a significant decrease in the overall responses of PVT^D2R+^ neurons during the reward approach compared to when they were tested while hungry (Fig. 1D). Additionally, trials conducted across the different ‘trial groups’ in the sated sessions showed significantly higher latencies to approach (Fig. 1E) and more pronounced decreases in PVT^D2R+^ neuronal activity (Fig. 1F-H). PVT^D2R+^ neuronal activity during sated states was also negatively associated with trial order (Fig. 1I,J), such that lower signals were associated with later trials, but not with latency to approach (Fig. 1K). Conversely, when the same mice were tested while hungry and given a sweet non-caloric reward, PVT^D2R+^ neuronal activity significantly increased during the reward approach (Fig. 1D). However, comparing early (G1) and late (G5) group trials in the H-NCal sessions showed similar response magnitudes (Fig. 1L,M). Additionally, the approach AUC was no longer significantly correlated with trial order (Fig. 1N,O) or latency (Fig. 1P). The lack of modulation by the physiological state during H-NCal sessions persisted even after mice were exposed to a non-caloric reward for 4 hours daily over two weeks, teaching them that non-caloric rewards do not satisfy them (Supp. Fig. 2).

Interestingly, in a new cohort of mice tested first while hungry using a caloric reward (H-SMS sessions) and a non-caloric reward (H-NCal sessions), followed by a sated non-caloric reward (S-NCal sessions; Fig. 1R), we found no differences in the magnitude of the PVT^D2R+^ neuronal photometric responses between these three distinct sessions (Fig. 1S-V). These findings show that even when mice are sated, a non-caloric reward still resulted in robust increases in PVT^D2R+^ neuronal activity during reward approach. Importantly, sated mice performed equivalent number of trials as hungry mice (Fig. 1Q,V), suggesting that the decreases in PVT^D2R+^ neuronal photometric responses during sated states are not solely due to an overall reduction in motivation. Altogether, these results strongly suggest that PVT^D2R+^ neurons can track the internal state of the animal. Additionally, these results also suggest that one way they track hunger levels is by sensing the caloric intake.

### PVT^D2R+^ neuronal activity is influenced by the interoceptive value of the outcome

Our findings showed that a non-caloric reward elicited the same response magnitude regardless of state, indicating PVT^D2R+^ neurons might signal the potential interoceptive value of the reward. Indeed, interoceptive value refers to the subjective value assigned to the internal physiological state or bodily sensations used to influence decision-making and behavior signals^2,4,9^. To explore this, we tested whether the interoceptive value of the outcome affected PVT^D2R+^ neuronal activity. We compared the responses of PVT^D2R+^ neurons during three separate sessions where we varied the relative and the interoceptive values of the outcomes (Fig. 2A). First, we examined the responses of PVT^D2R+^ neurons when hungry mice received SMS reward which has a ‘higher relative value’, 10% sucrose solution (Suc) which has a ‘medium relative value’, and plain water (H_2_O) which has a ‘low relative value’. We hypothesized that if PVT^D2R+^ neurons were sensitive to the relative value of the reward, switching from SMS to a lower value outcome would decrease the photometric signal during reward approach. However, we found that the response magnitudes during reward approach were similar between H-SMS and H-Suc (Fig. 2B). Trials within the H-Suc sessions also showed state-dependent effects (Fig. 2C-F), though these were less pronounced than with the calorie-dense SMS reward. Additionally, there was a negative association between PVT^D2R+^ neuronal signal and trial order in the H-Suc sessions (Fig. 2G,H). When comparing reward approach during sessions where hungry mice received water (H-H_2_O) as the outcome, we observed a significant decrease in PVT^D2R+^ neuronal responses (Fig. 2B). This decrease occurred quickly upon encountering the new outcome, with no significant differences in signal magnitude between early (G1) and late (G5) trial groups (Fig. 2I-K). In addition, there was no longer an association between signal magnitude and trial order during the approach (Fig. 2L), but the FLMM analysis revealed a negative association after reward zone entry (Fig. 2M). These decreases were likely not due to changes in overall motivation, as mice performed an equivalent number of trials across the test sessions (Fig. 2N). These findings suggest that PVT^D2R+^ neurons are activated by behaviors leading to physiologically relevant outcomes, likely encoding interoceptively-valuable stimuli rather than relative value. Indeed, in a 4-bottle choice test, mice strongly preferred the calorie-rich SMS when given the choice between SMS, 10% sucrose, a non-caloric reward, and water (Fig. 2O).

**Figure 2.**
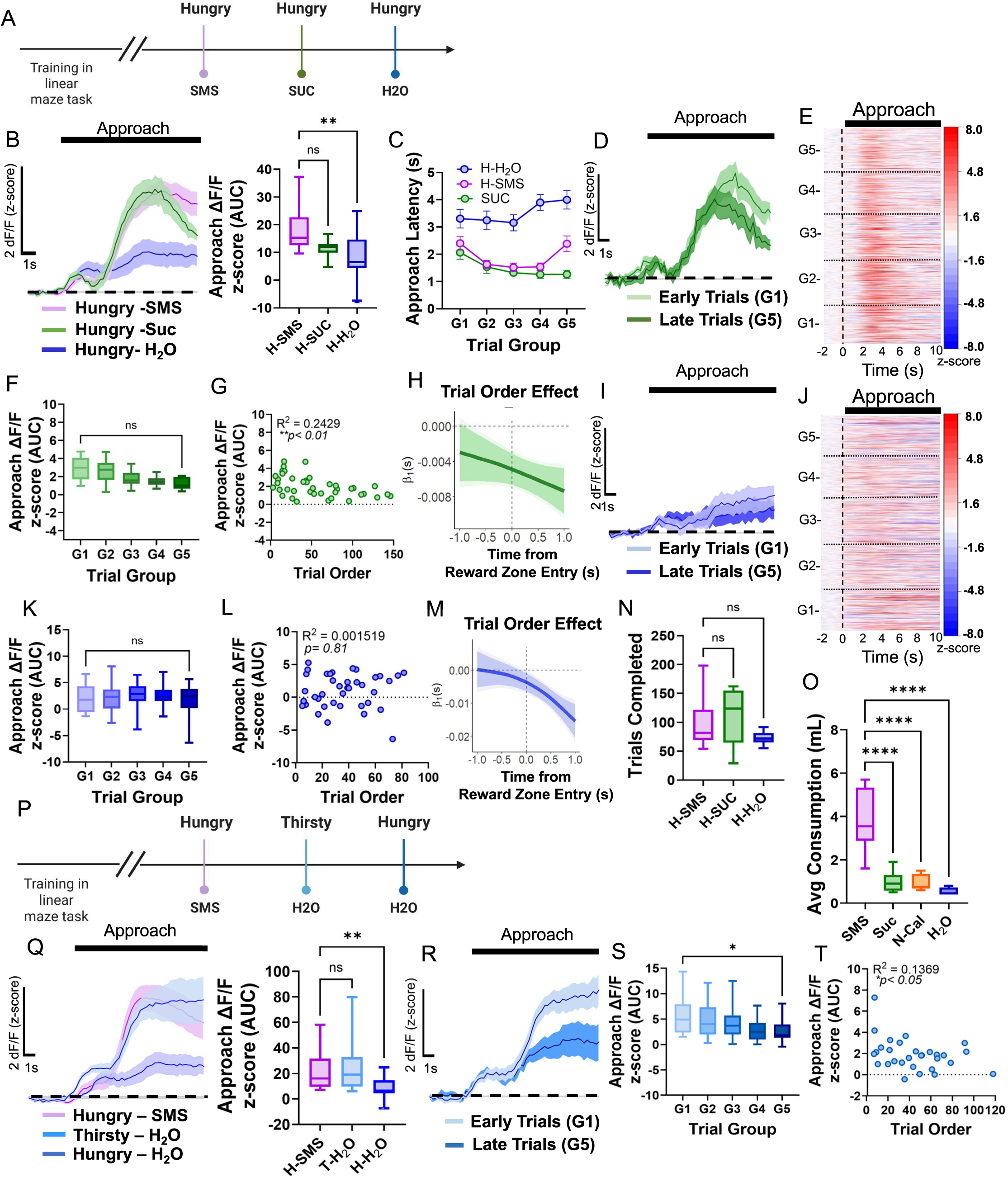
PVT^D2R+^ neuronal activity is modulated by the interoceptive value of the outcome. **(A)** Timeline schematic for the test sessions. **(B)** Average GCaMP6s responses (n=5) from PVT^D2R+^ neurons and AUC quantification comparing ramping activity during reward approach in test sessions: H-SMS: n = 460 trials, H-Suc: n = 885 trials and H-H_2_O: n = 549 trials. Repeated measures ANOVA, \*\**p<0.01*. H-SMS vs. H-Suc Dunnett’s multiple comparisons test *p=0.12, ns.* H-SMS vs. H-H_2_O Dunnett’s multiple comparisons test \*\**p<0.01*. **(C)** Latencies to reach the reward zone across ‘trial group blocks’ (G1 – G5) for H-SMS, H-Suc, and H-H_2_O sessions. Two-way ANOVA, Effect of trial group block *\*p<0.05*; Effect of test session \*\*\*\**p<0.0001;* Interaction *\*p<0.05*. **(D)** Average GCaMP6s responses from PVT^D2R+^ neurons comparing reward approach between early ‘G1’ and late ‘G5’ trials within the H-Suc test sessions. **(E)** Heatmap of reward approach responses from PVT^D2R+^ neurons during H-Suc test sessions, time-locked to approach onset, sorted by trial order, and binned into 5 ‘trial group blocks’ (G1 – G5). **(F)** AUC quantification of the reward approach-evoked changes in GCaMP6s activity across trial group blocks within the H-Suc test sessions. Repeated measures ANOVA, *\*p<0.05*. G1 vs. G5 Dunnett’s multiple comparisons test *p=0.05, ns.* **(G)** Correlation between the AUC of the reward approach-evoked changes in GCaMP6s activity and trial order within the H-Suc test sessions. **(H)** FLMM coefficient estimates plot of the approach trial order effect and statistical significance at each trial time-point results for the photometric responses of PVT^D2R+^ neurons for reward zone entry. The plot shows a negative association between PVT^D2R+^ GCaMP6s responses and trial order before and after reward zone entry for the H-Suc test sessions. **(I)** Average GCaMP6s responses from PVT^D2R+^ neurons comparing reward approach between early ‘G1’ and late ‘G5’ trials within the H-H_2_O test sessions. **(J)** Heatmap of reward approach responses from PVT^D2R+^ neurons during H-H_2_O test sessions. **(K)** AUC quantification of the reward approach-evoked changes in GCaMP6s activity across trial group blocks within the H-H_2_O test sessions. Repeated measures ANOVA, *p=0.50, ns*. **(L)** Correlation between the AUC of the reward approach-evoked changes in GCaMP6s activity and trial order within the H-H_2_O test sessions. **(M)** FLMM coefficient estimates plot of the approach trial order effect and statistical significance at each trial time-point results for the photometric responses of PVT^D2R+^ neurons for reward zone entry. The plot shows a negative association between PVT^D2R+^ GCaMP6s responses and trial order only after reward zone entry for the H-H_2_O test sessions. **(N)** Quantification of trials completed during H-SMS, H-Suc, and H-H_2_O test sessions. Repeated measures ANOVA, \**p<0.05.* H-SMS vs. H-Suc Dunnett’s multiple comparisons test *p=0.63, ns*. H-SMS vs. H-H_2_O Dunnett’s multiple comparisons test *p=0.06, ns*. **(O)** Quantification of average consumption (mLs) in a 4-bottle test of SMS, 10% Suc, Non-Cal and H_2_O *ad libitum* consumption. Repeated measures ANOVA, \*\*\*\**p<0.0001.* **(P)** Schematic of the timeline for the test sessions. **(Q)** Average GCaMP6s responses (n=7) from PVT^D2R+^ neurons and AUC quantification comparing ramping activity during reward approach in test sessions: H-SMS, T-H_2_O, and H-H_2_O. H-SMS: n = 1111 trials, T-H_2_O: n = 864 trials and H-H_2_O: n = 762 trials. Repeated measures ANOVA, \*\**p<0.01*. H-SMS vs. T-H_2_O Dunnett’s multiple comparisons test *p=0.97, ns.* H-SMS vs. H-H_2_O Dunnett’s multiple comparisons test \*\**p<0.01*. **(R)** Average GCaMP6s responses from PVT^D2R+^ neurons comparing reward approach between early ‘G1’ and late ‘G5’ trials within the T-H_2_O test sessions. **(S)** AUC quantification of the reward approach-evoked changes in GCaMP6s activity across trial group blocks within the T-H_2_O test sessions. Repeated measures ANOVA, \*\**p<0.01*. G1 vs. G5 Dunnett’s multiple comparisons test \**p<0.05*. **(T)** Correlation between the AUC of the reward approach-evoked changes in GCaMP6s activity and trial order within the T-H_2_O test sessions.

To further demonstrate that PVT^D2R+^ neurons encode physiologically relevant outcomes, we compared the mean photometric response magnitude of PVT^D2R+^ neurons across three conditions (Fig. 2P): 1) sessions where mice were tested while hungry and given the SMS reward (H-SMS), 2) sessions where mice were tested while hungry and given water (H-H_2_O), and 3) sessions where mice had *ad libitum* access to food, water was restricted, and water was given as a reward (T-H_2_O). We found that when mice were water-restricted and given water as a reward, there were robust increases in the photometric responses of PVT^D2R+^ neurons during reward approach. The magnitude of these responses was similar to those observed when mice were tested while hungry and given the SMS reward (Fig. 2Q). Importantly, we again observed state-dependent effects, with higher activity during early trials as compared to late trials (Fig. 2R,S), and a negative association between signal magnitude and trial order (Fig. 2T). Overall, these data strongly suggest that PVT^D2R+^ neurons are encoding the behaviors that lead to physiologically relevant outcomes.

### PVT^D2R+^ neuronal activity is state and outcome-dependent

Our findings showed that motivated behavior engages PVT^D2R+^ neurons only when the outcome is physiologically relevant, such as when hungry mice receive food or thirsty mice receive water. However, we previously observed that PVT^D2R+^ neurons remain active even when no reward is given during the linear task^39^. We hypothesized that this activity persists during reward omission due to the expectation of a reward. To test this, we trained mice for 8 weeks in rewarded sessions. This ensured that hungry mice maintained high response rates despite reward omission (H-OM), allowing us to test the effects of reward expectation on PVT^D2R+^ neuronal activity. We also included a session where sated mice experienced reward omission (S-OM) (Fig. 3A). We found that reward approach still elicited robust PVT^D2R+^ neuronal responses after extended training (H-SMS; Fig. 3B). However, complete reward omission (H-OM) significantly decreased PVT^D2R+^ neuronal responses during reward approach compared to H-SMS sessions (Fig. 3B) and increased the latency to approach (Fig. 3C). Additionally, state-dependent modulation of PVT^D2R+^ neurons was no longer observed (Fig. 3E,F), and there was no significant correlation between approach AUC and trial order (Fig. 3G). Surprisingly, FLMM analysis revealed a positive association between trial order and PVT^D2R+^ neuronal response magnitude (Fig. 3H). During the sated omission session (S-OM), PVT^D2R+^ neuronal responses were not significantly modulated by reward approach (Fig. 3B,I-M). Interestingly, mice in the H-OM session performed more trials than in the H-SMS sessions (Fig. 3N). These findings suggest that the decreases in signals were likely not solely due to a loss of motivation.

**Figure 3.**
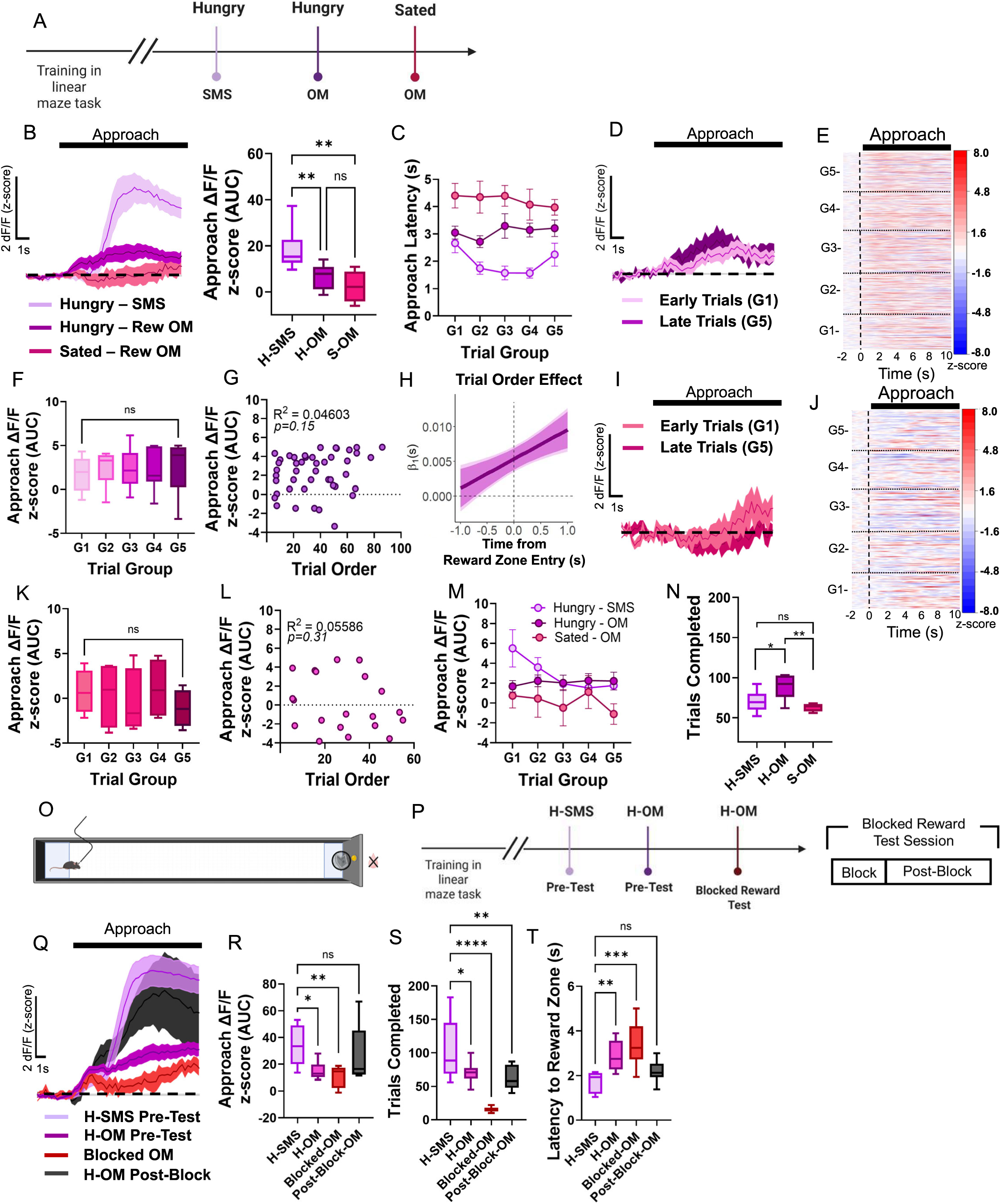
PVT^D2R+^ neuronal activity is state and outcome dependent. **(A)** Timeline schematic for the test sessions. **(B)** Average GCaMP6s responses (n=4) from PVT^D2R+^ neurons and AUC quantification comparing ramping activity during reward approach in test sessions: H-SMS, H-OM, and S-OM. H-SMS: n = 705 trials, H-OM: n= 797 trials and S-OM: n = 253 trials from 5 mice. Repeated measures ANOVA, \*\**p<0.01*. H-SMS vs. H-OM Tukey’s multiple comparisons test \*\**p<0.01*. H-SMS vs. S-OM Tukey’s multiple comparisons test \*\**p<0.01.* H-OM vs. S-OM Tukey’s multiple comparisons test *p=0.58, ns.* **(C)** Latencies to reach the reward zone across ‘trial group blocks’ (G1 – G5) for H-SMS, H-OM, and S-OM sessions. Two-way ANOVA, Effect of trial group block *p=0.54*, *ns.* Effect of test session \*\*\*\**p<0.0001;* Interaction *p=0.44, ns*. **(D)** Average GCaMP6s responses from PVT^D2R+^ neurons comparing reward approach between early ‘G1’ and late ‘G5’ trials within the H-OM test sessions. **(E)** Heatmap of reward approach responses from PVT^D2R+^ neurons during H-OM test sessions, time-locked to approach onset, sorted by trial order, and binned into 5 ‘trial group blocks’ (G1 – G5). **(F)** AUC quantification of the reward approach-evoked changes in GCaMP6s activity across trial group blocks within the H-OM test sessions. Repeated measures ANOVA, *p=0.45, ns*. **(G)** Correlation between the AUC of the reward approach-evoked changes in GCaMP6s activity and trial order within the H-OM test sessions. **(H)** FLMM coefficient estimates plot of the approach trial order effect and statistical significance at each trial time-point results for the photometric responses of PVT^D2R+^ neurons for reward zone entry. The plot shows a positive association between PVT^D2R+^ GCaMP6s responses and trial order before and after reward zone entry for the H-OM test sessions. **(I)** Average GCaMP6s responses from PVT^D2R+^ neurons comparing reward approach between early ‘G1’ and late ‘G5’ trials within the S-OM test sessions. **(J)** Heatmap showing reward approach responses from PVT^D2R+^ neurons during S-OM test sessions. **(K)** AUC quantification of the reward approach-evoked changes in GCaMP6s activity across trial group blocks within the S-OM test sessions. Repeated measures ANOVA, *p=0.45*, *ns.* **(L)** No correlation between the AUC of the reward approach-evoked changes in GCaMP6s activity and trial order within the S-OM test sessions. **(M)** AUC quantification of the reward approach across ‘trial group blocks’ (G1 – G5) for H-SMS, H-OM, and S-OM sessions. Two-way ANOVA, Effect of trial group block *p=0.26*; *ns.* Effect of test session \*\**p<0.01;* Interaction *p=0.27, ns*. **(N)** Quantification of trials completed within a session comparing trials performed during the H-SMS, the H-OM and the S-OM test sessions. Repeated measures ANOVA, \*\**p<0.01.* H-SMS vs. H-OM Tukey’s multiple comparisons test \**p<0.05*. H-SMS vs. S-OM Tukey’s multiple comparisons test *p=0.60, ns*. H-OM vs. S-OM Tukey’s multiple comparisons test \*\**p<0.01*. **(O)** Schematic illustrating the blocked reward test in the linear maze task. **(P)** *Left*: Timeline schematic for the blocked reward test sessions. *Right*: Schematic of the blocked reward test session showing that each session was divided into 2 parts in which the reward tray was blocked for the first 15 min of the session and was then unblocked for the remainder of the session. **(Q)** Average GCaMP6s responses from PVT^D2R+^ neurons (n=4) and **(R)** AUC quantification comparing ramping activity during reward approach for the H-SMS pre-test, H-OM pre-test, Blocked OM, and H-OM post-block tests sessions. Repeated measures ANOVA, \*\**p<0.01.* H-SMS pre-test vs. H-OM pre-test Tukey’s multiple comparisons test \**p<0.05*. H-SMS pre vs. Blocked OM Tukey’s multiple comparisons test \*\**p<0.01*. H-SMS pre vs. H-OM post-block Tukey’s multiple comparisons test *p=0.69, ns*. **(S)** Quantification of total trials completed for the H-SMS pre-test, H-OM pre-test, Blocked OM, and H-OM post-block tests sessions. Repeated measures ANOVA, \*\*\*\**p<0.0001.* H-SMS pre-test vs. H-OM pre-test Tukey’s multiple comparisons test \**p<0.05*. H-SMS pre-test vs. Blocked OM Tukey’s multiple comparisons test \*\*\*\**p<0.*0001. H-SMS pre-test vs. H-OM post-block Tukey’s multiple comparisons test \*\**p<0.001*. H-OM pre-test vs. H-OM post-block Tukey’s multiple comparisons test *p=0.92, ns*. **(T)** Quantification of latency to reward zone for the H-SMS pre-test, H-OM pre-test, Blocked OM, and H-OM post-block tests sessions. Repeated measures ANOVA, \*\*\**p<0.001.* H-SMS pre-test vs. H-OM pre-test Tukey’s multiple comparisons test \*\**p<0.01*. H-SMS pre-test vs. Blocked OM Tukey’s multiple comparisons test \*\*\**p<0.001*. H-SMS pre-test vs. H-OM post-block Tukey’s multiple comparisons test *p=0.52, ns*.

**Figure 4.**
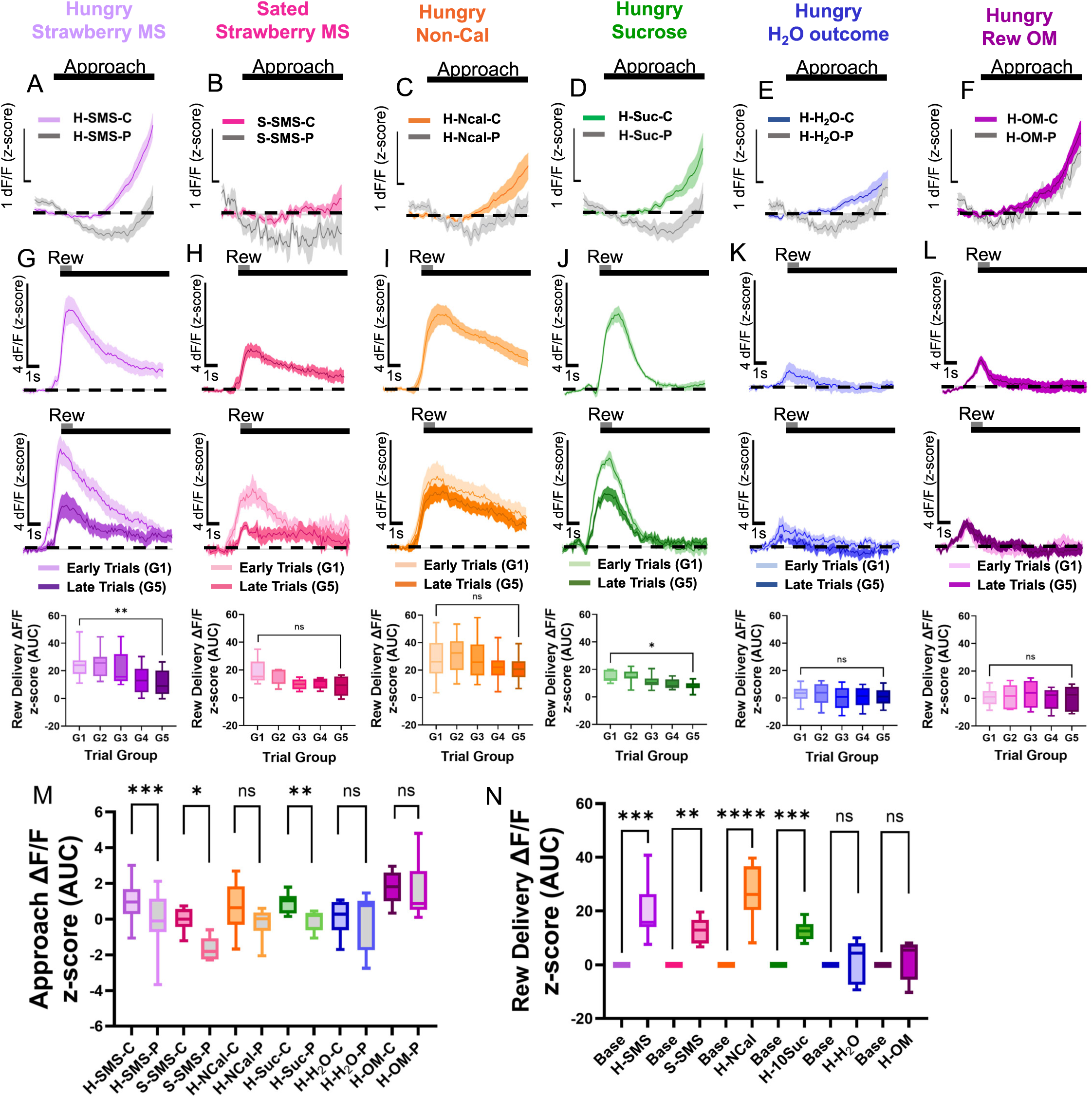
PVT^D2R+^ neuronal activity during approach only is modulated by outcome and outcome expectation. **(A-F)** Average GCaMP6s responses from PVT^D2R+^ neurons during approach only, normalized by linear interpolation comparing correct and premature trials in the test sessions: **(A)** H-SMS, **(B)** S-SMS, **(C)** H-NCal, **(D)** H-Suc, **(E)** H-H_2_O, and **(F)** H-OM**. (G-L)** Average GCaMP6s responses from PVT^D2R+^ neurons (top), comparisons between early ‘G1’ and late ‘G5’ trials (middle) and AUC quantifications across trial groups (bottom) during reward delivery for the **(G)** H-SMS test sessions; Repeated measures ANOVA, \*\*\*\**p<0.0001.* G1 vs. G5 Dunnett’s multiple comparisons test \*\**p<0.01*, **(H)** for S-SMS test sessions; Repeated measures ANOVA, *p=0.10, ns.* G1 vs. G5 Dunnett’s multiple comparisons test *p=0.08, ns*, **(I)** for H-NCal test sessions; Repeated measures ANOVA, *p=0.17, ns.* G1 vs. G5 Dunnett’s multiple comparisons test *p=0.65, ns*, **(J)** for H-Suc test sessions; Repeated measures ANOVA, \*\*\**p<0.001.* G1 vs. G5 Dunnett’s multiple comparisons test \**p<0.05*, **(K)** for H-H_2_O test sessions; Repeated measures ANOVA, *p=0.48, ns.* G1 vs. G5 Dunnett’s multiple comparisons test *p=0.87, ns*, **(L)** and for H-OM test sessions; Repeated measures ANOVA, *p=0.32, ns.* G1 vs. G5 Dunnett’s multiple comparisons test *p=0.98, ns*. **(M)** AUC quantifications of the reward approach only evoked changes in GCaMP6s activity across the different test sessions for correct ‘C’ and premature ‘P’ trials. Two-tailed paired t-tests. H-SMS-C vs. H-SMS-P: \*\*\**p<0.001.* S-SMS-C vs. S-SMS-P: \**p<0.05.* H-NCal-C vs. H-NCal-P: *p=0.07, ns.* H-Suc-C vs. H-Suc-P: \*\**p<0.01.* H-H_2_O-C vs. H-H_2_O-P *p=0.69, ns.* H-OM-C vs. H-OM-P *p=0.75, ns.* Two-tailed unpaired t-tests. H-SMS-C vs. H-OM-C \**p<0.05.* H-SMS-P vs. H-OM-P \*\**p<0.01.* **(N)** AUC quantifications of the reward delivery and feeding bouts-evoked changes in GCaMP6s activity across the different test sessions. Two-tailed paired t-tests. H-SMS baseline (base) vs. H-SMS: \*\*\**p<0.001.* Base vs. S-SMS: \*\**p<0.01.* Base vs. H-NCal: \*\*\*\**p<0.0001.* Base vs. H-Suc: \*\*\**p<0.001.* Base vs. H-H_2_O: *p=0.50, ns.* Base vs. H-OM: *p=0.60, ns*.

To further investigate how reward expectation affects PVT^D2R+^ neuronal activity during reward omission, we examined the neuronal responses when access to a reward tray was physically blocked (Fig. 3O-T), and whether removing this block (without delivering the reward) would increase reward expectation and thus enhance PVT^D2R+^ photometric responses. We hypothesized that a physical barrier serves as a cue signaling inaccessibility, thereby lowering reward expectation. Conversely, removing the barrier could raise expectation by altering the animal’s predictive model of reward availability. To test this, we first trained hungry mice in a standard SMS pre-test to establish baseline neuronal responses (Fig. 3P,Q). Next, we conducted an omission (OM) pre-test, confirming that reward omission reduced PVT^D2R+^ activity (Fig. 3P,Q). Finally, the same mice underwent a blocked reward test session (Fig. 3P), which had two phases: 1) *Block Phase* (15 minutes)— a wire mesh physically blocked access to the reward port, 2) *Post-Block Phase*— The mesh was removed, but the reward remained omitted. This design allowed us to assess whether the removal of the physical block alone could elevate reward expectation and modulate PVT^D2R+^ activity. We found that physically blocking the reward port significantly reduced PVT^D2R+^ photometric responses (Fig. 3Q,R). Notably, once the physical block was removed (despite the continued absence of the reward) the photometric responses significantly increased, reaching levels comparable to those observed during the H-SMS pre-test (Fig. 3Q,R).

Next, to further test the influence of outcome expectation on PVT^D2R+^ neuronal activity, we also analyzed the photometric responses of PVT^D2R+^ neurons during ‘premature’ trials performed by hungry mice. In the linear maze task, mice had to wait for 2 seconds in the trigger zone to receive the cue signaling reward availability. However, in some trials, mice did not wait properly in the trigger zone before approaching, resulting in the absence of the cue during the reward approach. These trials were categorized as’premature approaches’. We found that during premature approaches, mean PVT^D2R+^ neuronal activity was significantly lower during the approach to the reward zone (Supp. Fig. 3A). Importantly, the latencies to approach were equal between the “correct” rewarded approaches and unrewarded premature approaches (Supp. Fig. 3B). Interestingly, we observed state-dependent effects, with higher activity during early trials (G1) compared to late trials (G5) in the premature trials (Supp. Fig. 3C,D). Together, these findings suggest that PVT^D2R+^ neuronal activity is modulated within a session depending on outcome expectation.

Lastly, we aimed to test the hypothesis that PVT^D2R+^ neurons are responding to two dissociable but related events: 1) the expectation of an outcome during the approach phase, and 2) the outcome itself. However, because our behavioral task is entirely self-paced, the duration of each trial varies. This variability in trial latencies made it difficult to cleanly separate the neural responses associated with reward expectation during approach from those associated with reward delivery when averaging across grouped trials. To address this limitation and better dissociate PVT^D2R+^ neuronal responses to ‘approach only’ versus ‘reward only’ events, we conducted two complementary analyses. In the first analysis, we restricted each trial to either the approach phase or the reward delivery phase. We then calculated the AUC for each approach-only trial within each outcome group (see methods). This approach allowed us to independently examine neural activity during approach and during reward delivery. During the approach phase, we observed that in most test sessions, PVT^D2R+^ neuronal responses were significantly higher in correct trials compared to premature trials (Fig. 4A–F, M). Furthermore, the magnitude of these responses was modulated by the expected value of the outcome: sessions involving higher-value outcomes (e.g., H-SMS) elicited stronger PVT^D2R+^ activity than those with lower-value outcomes (Fig. 4A–F). Interestingly, in H-OM sessions, we found no significant difference in PVT^D2R+^ responses between correct and premature trials (Fig. 4F, M). This lack of differentiation is likely due to the ambiguity introduced by the absence of any outcome in both trial types during H-OM sessions. Unlike blocked access, which directly cued the inaccessibility of reward (Fig. 3O-T), the animal may be unable to determine whether the task contingencies have changed or whether the outcome was simply omitted, leading to a disruption in expectation. In the second analysis, we isolated and compared PVT^D2R+^ neuronal responses specifically during reward delivery and feeding bouts (the first 10 sec after reward delivery). We found that these photometric responses were also influenced by both the nature of the outcome and the internal state of the mice (Fig. 4G–L). Notably, we observed significant state-dependent changes in PVT^D2R+^ activity within sessions that included caloric rewards (Fig. 4G, H, J, N). These state-dependent modulations were absent in sessions that did not involve caloric rewards, such as H-NCal (Fig. 4I, N), H-H_2_O (Fig. 4E, N), and H-OM (Fig. 4F, N).

In our second analysis, we applied a concurrent functional linear mixed model (FLMM), which allows for comparisons across sessions while accounting for variability in trial durations. This model included a binary functional covariate that marked timepoints when the animal was approaching the reward versus when it was receiving the reward. This approach enabled us to examine how behavioral phase (approach vs. outcome delivery), internal state (e.g., hunger), and outcome type (e.g., caloric vs. non-caloric) dynamically influence PVT^D2R+^ photometric responses. The concurrent FLMM analysis confirmed our previous findings and revealed additional differences that were not detectable using trial-averaged methods (Supp. Fig. 4). Specifically, during reward delivery, the mean photometry signal of PVT^D2R+^ neurons was significantly higher in the H-SMS condition compared to S-SMS (Supp. Fig. 4A β_₁_*(s)*), H-H_₂_O (Supp. Fig. 4D β_₁_*(s)*), and H-OM (Supp. Fig. 4E β_₁_*(s)*). However, in the H-NCal condition, the mean photometry signal was significantly higher than H-SMS during the later stages of reward delivery (Supp. Fig. 4B β *(s)*), suggesting that PVT^D2R+^ neurons respond similarly to caloric and non-caloric rewards, but only caloric rewards induce state-dependent decreases during prolonged feeding. No significant differences were observed between H-SMS and H-Suc (Supp. Fig. 4C β (s)), indicating that PVT^D2R+^ neurons do not encode relative reward value. Across all conditions, the model showed that PVT^D2R+^ mean signal was consistently lower during approach than during reward delivery (Supp. Fig. 4A–E β_₂_*(s)*). Lastly, this FLMM model enabled us to evaluate the interaction effects between behavioral states (e.g., approaching vs. receiving), internal states (e.g., hungry vs. sated), and outcome types (e.g., SMS vs. NCal), and how these factors collectively influence the neural response of PVT^D2R+^ neurons (Supp. Fig. 4A-F β*3(s)*).

Specifically, we observed that the interaction for the H-SMS vs. S-SMS comparison is negative, indicating that the mean differences in photometric responses between these two outcome types are larger during the outcome delivery than during outcome approach. The results were similar for the interaction in the H-SMS vs. H-H_2_O comparisons (Supp. Fig. 4D β*_3_(s)*) and the H-SMS vs. H-OM comparisons (Supp. Fig. 4E β*_3_(s)*). For the H-SMS vs. NCal comparison, we found that the interaction is first negative and then is positive, showing that the differences in mean neural activity between approach and reward delivery timepoints are lower in H-NCal (than H-SMS) during early portions of the trial. Then, during later portions of the trial, the difference in mean signals between approach and reward timepoints is higher in the H-SMS condition (Supp. Fig. 4B β*_3_(s)*). Finally, we found no differences between H-SMS and H-Suc (Supp. Fig. 4C β*_3_(s)*). Overall, the concurrent FLMM analysis demonstrates that PVT^D2R+^ photometric signals are shaped by a combination of the behavioral phase, expected outcome, and internal state.

### PVT^D2R+^ neuronal activity during return is modulated by state, outcome, and outcome expectation

Next, we sought to test the *in vivo* dynamics of these PVT^D2R+^ neurons during the return from the reward zone to the trigger zone. We reasoned that by examining the *in vivo* dynamics during this time, where mice are engaging in the same motivated behavior (running) but not expecting a reward, would allow us to further dissociate the PVT^D2R+^ neuronal responses between expectation of the outcome and the outcome delivery. Interestingly, we found returning from the reward zone to the trigger zone resulted in a robust decrease in the photometric responses of PVT^D2R+^ neurons in all outcome sessions (Fig. 5). Moreover, the responses were also robustly modulated by the state of the mice, the outcome delivered, and the outcome expectation (Fig. 5A-H). Surprisingly, our analysis revealed state-dependent differences when comparing early trials (’G1’) to late trials (’G5’) in sessions where outcomes that led to greater satiation were provided (Fig. 5A,B,D,F). Specifically, stronger decreases in the signal of PVT^D2R+^ neurons were observed in early trials as compared to late trials. These effects again were absent in those sessions where non-caloric outcomes were provided (H-NCal, H-H_2_O, and H-OM sessions; Fig. 5C,E,G). Importantly, the decreases observed during return and the differences in modulation across various conditions (Fig. 5H) were not due to activity levels returning to baseline after reward delivery. First, mice spent ∼20 seconds in the reward zone before returning (Supp. Fig. 5), allowing reward-evoked signals to return to baseline in most groups (Fig. 4G–L). Second, trials with reward delivery and return responses were aligned and analyzed based on reward zone arrival and sorted by the time spent in the reward zone. These trials showed that the initiation of return closely aligned with decreases in the signal of PVT^D2R+^ neurons (Fig. 5I-L). Additionally, injections of the GABA sensor (GABASnFR2) into the PVT revealed increased GABA signaling during return (Supp. Fig. 6A-G), indicating that the decreases during the return phase are due to increases in inhibitory inputs. Interestingly, GABAergic inhibitory signaling remained consistent across trial groups (Supp. Fig. 6H-O), indicating that the observed modulations during return are unlikely due to changes in inhibitory signaling onto these neurons. Further analysis showed that PVT^D2R+^ neuronal responses increased immediately prior to return initiation (Supp. Fig. 7A; Fig. 5I-L). Moreover, the magnitude of these peaks in the signal during the decision to return drove the observed modulation (Supp. Fig. 7B-J). These findings suggest that PVT^D2R+^ neuronal activity increases during the decision to return, which is modulated by internal state, outcome, and expectation, and is then suppressed by inhibitory signaling during return onset.

**Figure 5.**
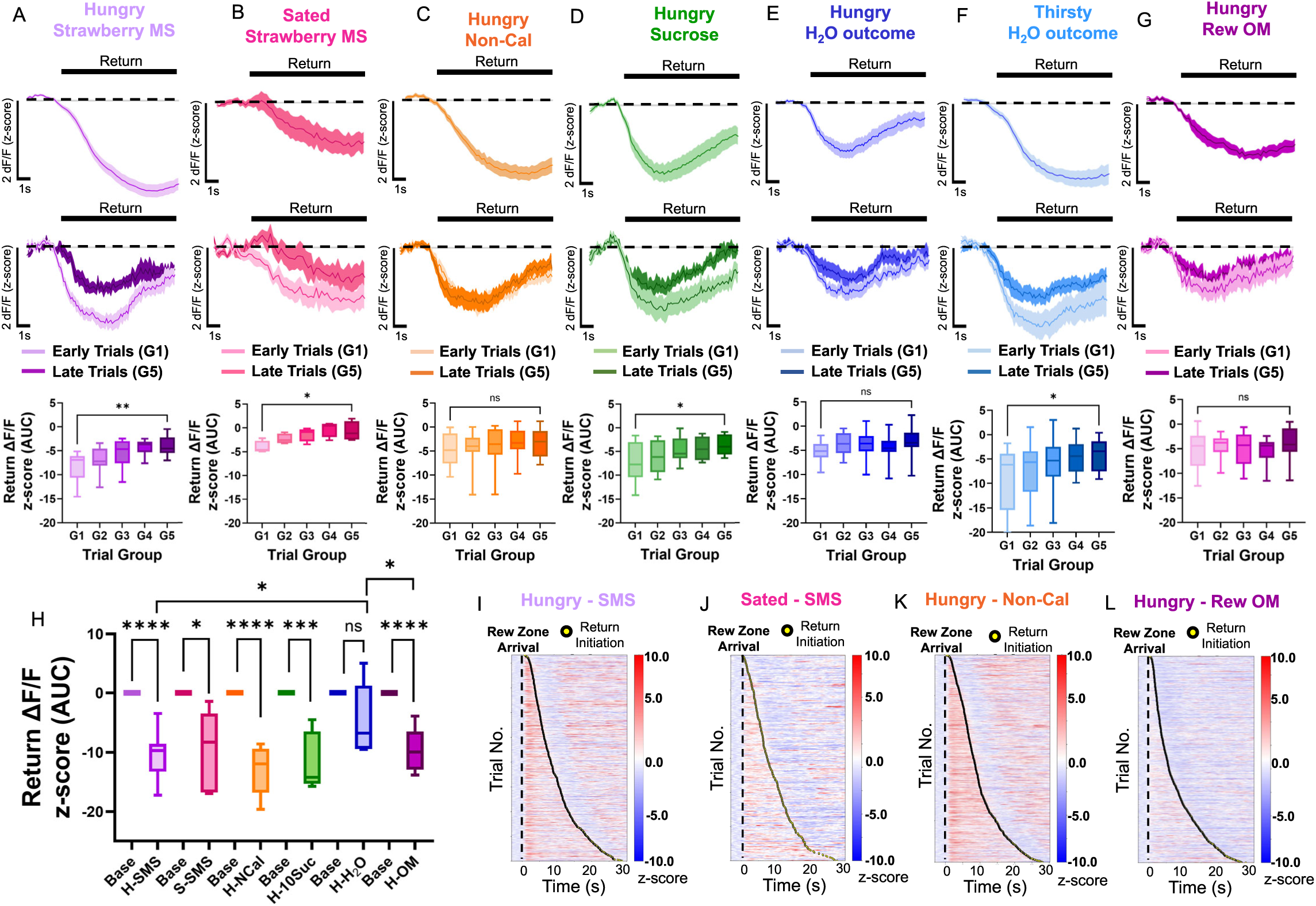
PVT^D2R+^ neuronal activity during return is modulated by outcome and outcome expectation. **(A-F)** Average GCaMP6s responses from PVT^D2R+^ neurons (top), comparisons between early ‘G1’ and late ‘G5’ trials (middle), and AUC quantifications across trial groups (bottom). These responses are time-locked to return onset, sorted by trial order, and binned into 5 trial group blocks (G1 – G5) during return to the trigger zone for **(A)** H-SMS test sessions: Repeated measures ANOVA, \*\*\*\**p<0.0001.* G1 vs. G5 Dunnett’s multiple comparisons test \*\**p<0.01*; **(B)** for S-SMS test sessions: Repeated measures ANOVA, \*\**p<0.01.* G1 vs. G5 Dunnett’s multiple comparisons test \**p<0.05*; **(C)** for H-NCal test sessions: Repeated measures ANOVA, *p=0.06, ns.* G1 vs. G5 Dunnett’s multiple comparisons test *p=0.06, ns*; **(D)** for H-Suc test sessions: Repeated measures ANOVA, \**p<0.05.* G1 vs. G5 Dunnett’s multiple comparisons test \**p<0.05*; **(E)** for H-H_2_O test sessions: Repeated measures ANOVA, *p=0.09, ns.* G1 vs. G5 Dunnett’s multiple comparisons test *p=0.07, ns*; **(F)** for T-H_2_O test sessions: Repeated measures ANOVA, \*\**p<0.01.* G1 vs. G5 Dunnett’s multiple comparisons test \**p<0.05, ns*; **(G)** and for H-OM test sessions: Repeated measures ANOVA, *p=0.41, ns.* G1 vs. G5 Dunnett’s multiple comparisons test *p=0.63, ns*. **(H)** AUC quantifications of the return-evoked changes in GCaMP6s activity across the different test sessions. Two-tailed paired t-tests. H-SMS baseline (base) vs. H-SMS: \*\*\*\**p<0.0001.* Base vs. S-SMS: \**p<0.05.* Base vs. H-NCal: \*\*\*\**p<0.0001.* Base vs. H-Suc: \*\*\**p<0.001.* Base vs. H-H_2_O: *p=0.08, ns.* Base vs. H-OM: \*\*\*\**p<0.0001.* Two-tailed unpaired t-tests. H-SMS vs H-H_2_O: \**p<0.05.* H-H_2_O vs. H-OM: \**p<0.05.* **(I-L)** Heatmap of trials containing reward delivery and return responses from PVT^D2R+^ neurons, time-locked to reward zone arrival, sorted by duration in reward zone. Yellow dots indicate return onset for test sessions: **(I)** H-SMS (strong responses modulated by state), **(J)** S-SMS (weak responses modulated by state), **(K)** H-NCal (strong responses not modulated by state) and **(L)** H-OM (weak responses not modulated by state).

### Motivational states are associated with PVT^D2R+^ neuronal responses across different conditions

Our findings showed that PVT^D2R+^ neurons mediate interoceptive predictions by integrating information about the animal’s state and the expected outcome. Previously, we demonstrated that PVT^D2R+^ neuronal activity was closely linked to the latency to approach the reward zone in a linear maze task^39^. Based on these findings, we hypothesized that PVT^D2R+^ neuronal activity is associated with latency to approach because latency reflects the motivational strength of the interoceptive prediction. Specifically, lower latencies (fast trials) would be associated with stronger predictions and thus higher PVT^D2R+^ neuronal responses, while higher latencies (slow trials) would be associated with weaker predictions and thus attenuated PVT^D2R+^ neuronal responses. To test this, we sorted trials by approach latency (labeling fast trials as’L1’ and slow trials as’L5’; Supp. Fig. 8) and calculated the approach slope under various conditions. Consistent with our prior work, we found significant correlations between latency and slope in most conditions (Supp. Fig. 8A–D, F), with fast trials showing steeper slopes. However, this relationship was absent in the H-H_2_O (Supp. Fig. 8E) and S-OM (Supp. Fig. 8G) conditions. These results suggest that PVT^D2R+^ activity tracks the motivational strength of interoceptive predictions, modulated by state, outcome, and expectation.

### PVT^D2R+^ neurons detect changes in reward magnitude

Interoception models have posited that brain areas encoding interoceptive predictions have the capability to detect changes in reward magnitude^4–6,9,43^. Since interoceptive predictions help anticipate and regulate bodily conditions, they inherently detect changes in reward magnitude which allow the brain to efficiently allocate effort and attention to optimizing resource acquisition^4,43^. As such, we next tested whether PVT^D2R+^ neurons could detect changes in reward magnitude. To investigate this, we first trained hungry mice to perform the linear maze task to obtain a non-caloric reward (Fig. 6A). We used a non-caloric reward to avoid state-dependent changes that could confound our results. Once we confirmed this during a pre-test (Fig. 6B, E), the mice underwent a’reward magnitude test session’ consisting of three within-session trial blocks, each offering different reward sizes (small: 1µl, medium: 3µl, and large: 6µl; Fig. 6A). We found that PVT^D2R+^ photometric responses were modulated by the different reward magnitudes, such that bigger rewards resulted in higher response signaling of these neurons, while smaller reward magnitudes resulted in lower response magnitudes (Fig. 6C). No significant changes in approach latency were found between reward magnitude trial blocks (Fig. 6D). Notably, variations in reward magnitude did not alter photometric responses during the return phase (Fig. 6F), although return latencies were reduced (Fig. 6G). Altogether, these findings strongly suggest that PVT^D2R+^ neurons can detect changes in reward magnitude.

**Figure 6.**
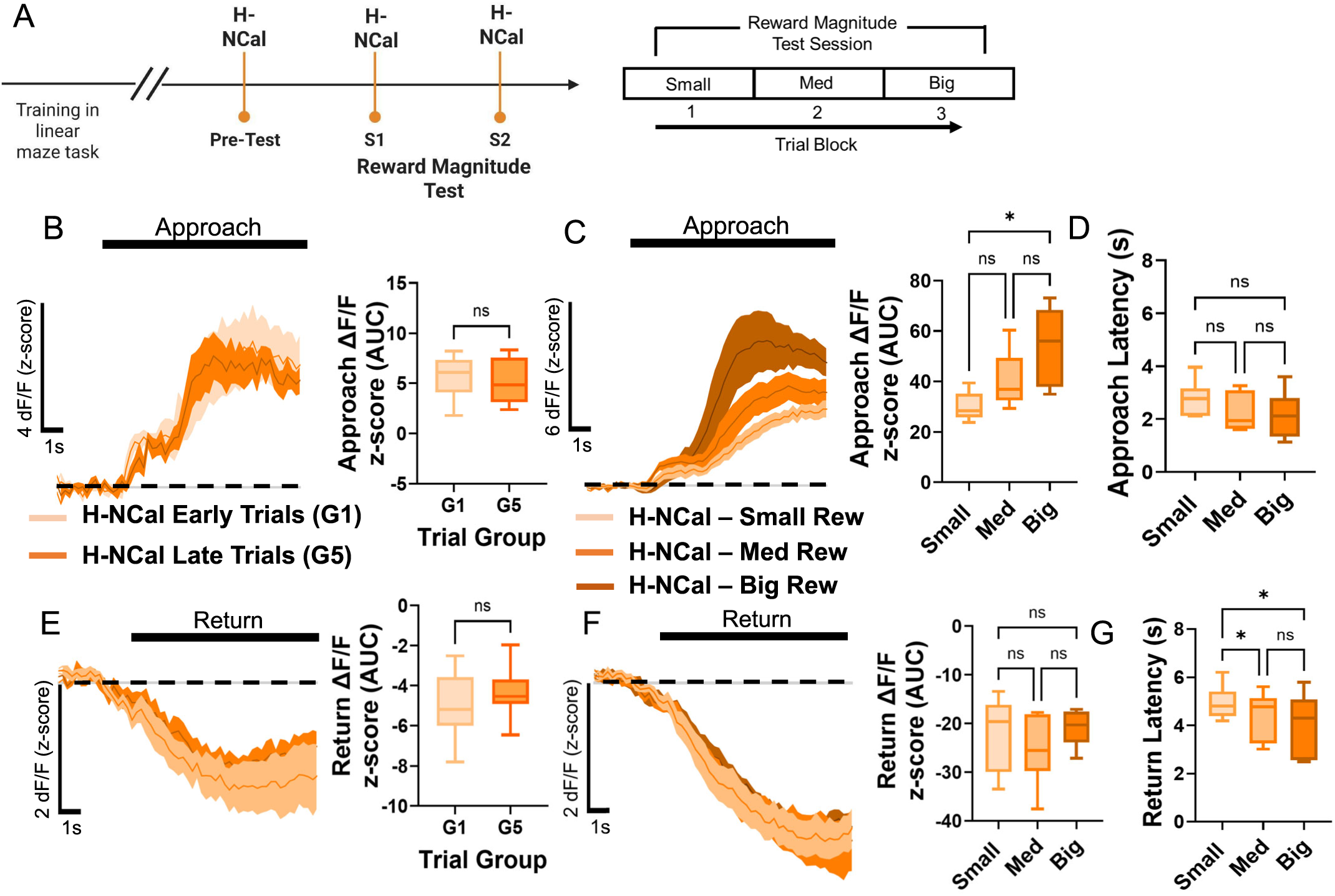
PVT^D2R+^ neurons detect changes in reward magnitude. **(A)** *Left:* Schematic timeline for test sessions measuring changes in reward magnitude. *Right:* Schematic of the reward magnitude test session showing that each session was divided into 3 trials blocks in which mice received NCal reward of different magnitudes: small (1µl), medium (3µl), and big (6µl). **(B)** *Left:* Average GCaMP6s responses (n=4) from PVT^D2R+^ neurons during the pre-test with a non-caloric reward. *Right:* AUC quantification comparing ramping activity during reward approach for early (G1) and late (G5) trials. Two-tailed paired t-test. G1 vs. G5, *p=0.48, ns.* **(C)** *Left:* Average GCaMP6s responses from PVT^D2R+^ neurons to the different reward amounts during approach. *Right:* AUC quantification comparing ramping activity during reward approach for the small, medium, and big reward. Repeated measures ANOVA, \**p<0.05.* Small vs. Medium: Tukey’s multiple comparisons test *p=0.27, ns*. Small vs. Big: Tukey’s multiple comparisons test \**p<0.05*. Medium vs. Big: Tukey’s multiple comparisons test *p=0.16, ns*. **(D)** Latencies to reach the reward zone for the small, medium and big rewards. Repeated measures ANOVA, *p=0.06, ns.* Small vs. Medium: Tukey’s multiple comparisons test *p=0.06, ns*. Small vs. Big: Tukey’s multiple comparisons test *p=0.18, ns*. Medium vs. Big: Tukey’s multiple comparisons test *p=0.38, ns*. **(E)** *Left:* Average GCaMP6s responses from PVT^D2R+^ neurons during return phase of the pre-test session. *Right:* AUC quantification comparing ramping activity during reward approach for early (G1) and late (G5) trials. Two-tailed paired t-test. G1 vs. G5, *p=0.21, ns.* **(F)** *Left:* Average GCaMP6s responses from PVT^D2R+^ neurons during return. *Right:* AUC quantification showing comparisons between return activity for the small, medium, and big reward. Repeated measures ANOVA, *p=0.47, ns.* Small vs. Medium: Tukey’s multiple comparisons test *p=0.59, ns*. Small vs. Big: Tukey’s multiple comparisons test *p=0.96, ns*. Medium vs. Big: Tukey’s multiple comparisons test *p=0.48, ns*. **(G)** Latencies for the return for the small, medium and big rewards. Repeated measures ANOVA, \**p<0.05.* Small vs. Medium: Tukey’s multiple comparisons test \**p<0.05*. Small vs. Big: Tukey’s multiple comparisons test \**p<0.05*. Medium vs. Big: Tukey’s multiple comparisons test *p=0.33, ns*.

### PVT^D2R+^ neuronal responses emerge from learned cue-outcome associations

Brain areas involved in interoceptive predictions also have the distinct ability to predict internal states based on the current state, expected outcomes, and learned environmental cues^4–6,9^. Thus, we next tested whether PVT^D2R+^ neuronal responses also depend on the environmental cues associated with the outcome and state. Indeed, we showed that when mice returned from the reward zone to the trigger zone in the maze task, the photometric responses of PVT^D2R+^ neurons displayed robust decreases that were state-and outcome-dependent (Fig. 5). Thus, we hypothesized that the differences in PVT^D2R+^ neuronal responses between the reward approach and the return to the trigger zone could be due to the learned association between a side of the maze and the outcome associated with it (reward zone = reward vs. trigger zone = no reward). If the increases in PVT^D2R+^ neuronal activity during the approach, and decreases during the return, are due to the learned cue-outcome associations, then these would appear gradually as mice learn to associate each side of the maze with their respective outcome. To test this, we assessed the photometric responses of PVT^D2R+^ neurons as mice learned to perform our linear maze task. Here, we first exposed mice to the SMS and ensured mice were consuming it reliably (Fig. 7A). We found a small modulation of PVT^D2R+^ neuronal responses during the reward approach (Fig. 7B; Supp. Fig. 9A) and during the return to the trigger zone (Fig. 7E; Supp. Fig. 9B) throughout the first training session (S1). However, as mice received more training sessions and learned to associate the reward zone with the delivery of the SMS and the return with the lack of reward, the PVT^D2R+^ neuronal responses during approach (Fig. 7B; Supp. Fig. 9A) and during return emerged (Fig. 7E; Supp. Fig. 9B) in a training-and session-dependent manner. To identify precisely when responses emerged, we compared the dynamics between the early trial group (G1) versus the late trial group (G5) in training session 1. Surprisingly, no differences were found between them (Fig. 7H,J). We then analyzed training session 2 (Fig. 7I,K) and compared the dynamics between the early (G1) versus late trial group (G5). Again, we found small increases in the overall activity of the PVT^D2R+^ neurons that were present even in the early trial group (G1). These increases were slightly bigger in late trials of both sessions (Fig. 7H, I), however these were not statistically significant. Notably, the lack of within-session (‘G1’ vs. ‘G5’) statistical differences for S1 and S2 sessions is likely due to responses being occluded by the satiety effects present, particularly in later trials. Nonetheless, these findings show that these neuronal responses emerge slowly despite successful task acquisition, as evidenced by the increases in trials performed during later sessions (Fig. 7D). For the return responses, we found similar dynamics in the emergence of the responses (Fig. 7J,K). However, by session 2, we found a significant difference between early and late trials (Fig. 7K), presumably due to the certainty of not receiving a reward after return. Altogether, these findings show that PVT^D2R+^ neuronal responses develop as mice learn cue-outcome associations. These findings are also in accordance with the new proposed role of PVT^D2R+^ neurons in the signaling of interoceptive predictions. Indeed, it has been suggested that predictive interoceptive responses develop more slowly than exteroceptive predictions^4^ because interoceptive signals tend to be subtler and more diffused than exteroceptive signals^4^.

**Figure 7.**
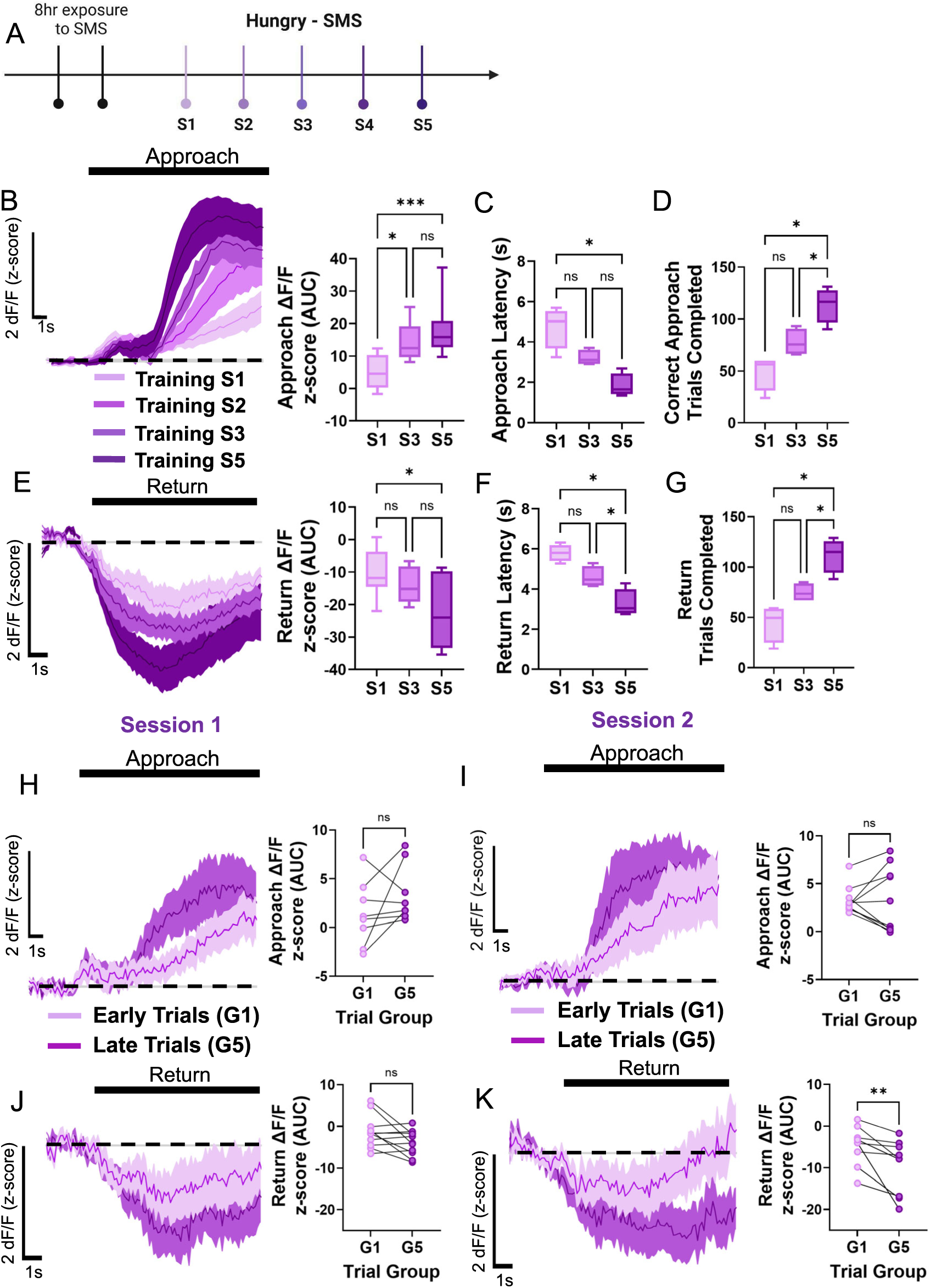
PVT^D2R+^ neuronal responses during approach and return emerge from the learning of cue–outcome associations. **(A)** Timeline schematic for the test sessions. **(B)** Average GCaMP6s responses (n=8) from PVT^D2R+^ neurons and AUC quantification comparing ramping activity during reward approach for the first ‘S1’, second ‘S2’, third ‘S3’ and fifth ‘S5’ training session in our linear maze task. Repeated measures ANOVA, \*\*\**p<0.001.* S1 vs. S3 Tukey’s multiple comparisons test \**p<0.05*. S1 vs. S5 Tukey’s multiple comparisons test \*\*\**p<0.001*. S3 vs. S5 Tukey’s multiple comparisons test *p=0.60, ns*. **(C)** Latencies to reach the reward zone for training sessions ‘S1’, ‘S3’ and ‘S5’. Repeated measures ANOVA, \*\**p<0.01.* S1 vs. S3 Tukey’s multiple comparisons test *p=0.06, ns*. S1 vs. S5 Tukey’s multiple comparisons test \**p<0.05*. S3 vs. S5 Tukey’s multiple comparisons test *p=0.08, ns*. **(D)** Correct approach trials performed during training sessions ‘S1’, ‘S3’ and ‘S5’. Repeated measures ANOVA, \**p<0.05.* S1 vs. S3 Tukey’s multiple comparisons test *p=0.25, ns*. S1 vs. S5 Tukey’s multiple comparisons test \**p<0.05*. S3 vs. S5 Tukey’s multiple comparisons test \**p<0.05.* **(E)** Average GCaMP6s responses from PVT^D2R+^ neurons and AUC quantification during return to trigger zone for the first ‘S1’, third ‘S3’, and fifth ‘S5’ training session. Repeated measures ANOVA, \**p<0.05.* S1 vs. S3 Tukey’s multiple comparisons test *p=0.54, ns.* S1 vs. S5 Tukey’s multiple comparisons test \**p<0.01*. S3 vs. S5 Tukey’s multiple comparisons test *p=0.14, ns*. **(F)** Latencies to return to the trigger zone for training sessions ‘S1’, ‘S3’, and ‘S5’. Repeated measures ANOVA, \**p<0.05.* S1 vs. S3 Tukey’s multiple comparisons test *p=0.15, ns*. S1 vs. S5 Tukey’s multiple comparisons test \**p<0.05*. S3 vs. S5 Tukey’s multiple comparisons test \**p<0.05*. **(G)** Total return trials performed during training sessions ‘S1’, ‘S3’, and ‘S5’. Repeated measures ANOVA, \**p<0.05.* S1 vs. S3 Tukey’s multiple comparisons test *p=0.25, ns*. S1 vs. S5 Tukey’s multiple comparisons test \**p<0.05*. S3 vs. S5 Tukey’s multiple comparisons test \**p<0.05.* **(H)** Average GCaMP6s responses and AUC quantification from PVT^D2R+^ neurons comparing reward approach between early ‘G1’ and late ‘G5’ trials for the first training session. Two-tailed paired t-test. G1 vs. G5, *p=0.31, ns.* **(I)** Average GCaMP6s responses and AUC quantification from PVT^D2R+^ neurons comparing reward approach between early ‘G1’ and late ‘G5’ trials for the second training session. Two-tailed paired t-test. G1 vs. G5, *p=0.91, ns.* **(J)** Average GCaMP6s responses from PVT^D2R+^ neurons comparing return to trigger zone between early ‘G1’ and late ‘G5’ trials for the first training session. Two-tailed paired t-test. G1 vs. G5, *p=0.05, ns.* **(K)** Average GCaMP6s responses from PVT^D2R+^ neurons comparing return to trigger zone between early ‘G1’ and late ‘G5’ trials for the second training session. Two-tailed paired t-tests. G1 vs. G5, \*\**p<0.01*.

### Chronic PVT^D2R+^ neuronal inhibition impairs performance in the linear maze tasks

Since PVT^D2R+^ neurons encode interoceptive predictions and their responses emerge slowly, across several sessions, we then aimed to investigate whether chronic chemogenetic inhibition of PVT^D2R+^ neurons would impair performance in our linear maze task. To do this, D2-Cre mice were injected with either an AAV-hSyn-DIO-hM4D(Gi)-mCherry virus or a control virus (AAV-CAG-Flex-TdTomato) in the PVT. Once the virus was expressed, hM4Di-PVT^D2R+^ mice and control-PVT^D2R+^ mice were injected with deschloroclozapine (DCZ; Fig. 8A,B) or CNO (Supp. Fig. 10) 30 mins prior to training in the linear maze task. Given that PVT^D2R+^ neuronal responses were slow to emerge (Supp. Fig. 9), we repeated our chemogenetic inhibition for three consecutive test sessions, followed by three sessions in which the mice were injected with saline prior to testing (Fig. 8B). We found that the chemogenetic inhibition of PVT^D2R+^ neurons did not result in acute changes in performance for both correct (Fig. 8C) and premature trials (Fig. 8E). We found, however, significant differences in the latencies to perform correct trials during the S2 and S3 (Fig. 8D), but no changes in latency to perform premature trials (Fig. 8F). Furthermore, we found that when mice were tested for post-inhibition effects, control mice performed a significantly higher number of correct trials compared to hM4Di-PVT^D2R+^ mice (Fig. 8C). No changes in the performance of premature trials were observed between controls and hM4Di-PVT^D2R+^ mice during post-inhibition testing (Fig. 8E). Lastly, we found that chemogenetic inhibition of PVT^D2R+^ neurons after task acquisition did not affect task performance (Supplementary Fig. 10 I,J). This suggests that these neurons are not essential for the execution or maintenance of task performance after the learning phase. Collectively, these results indicate that PVT^D2R+^ neurons play a critical role in early learning. Furthermore, prolonged inhibition of these neurons during training impairs the sustained performance of behaviors guided by interoceptive motivation.

**Figure 8.**
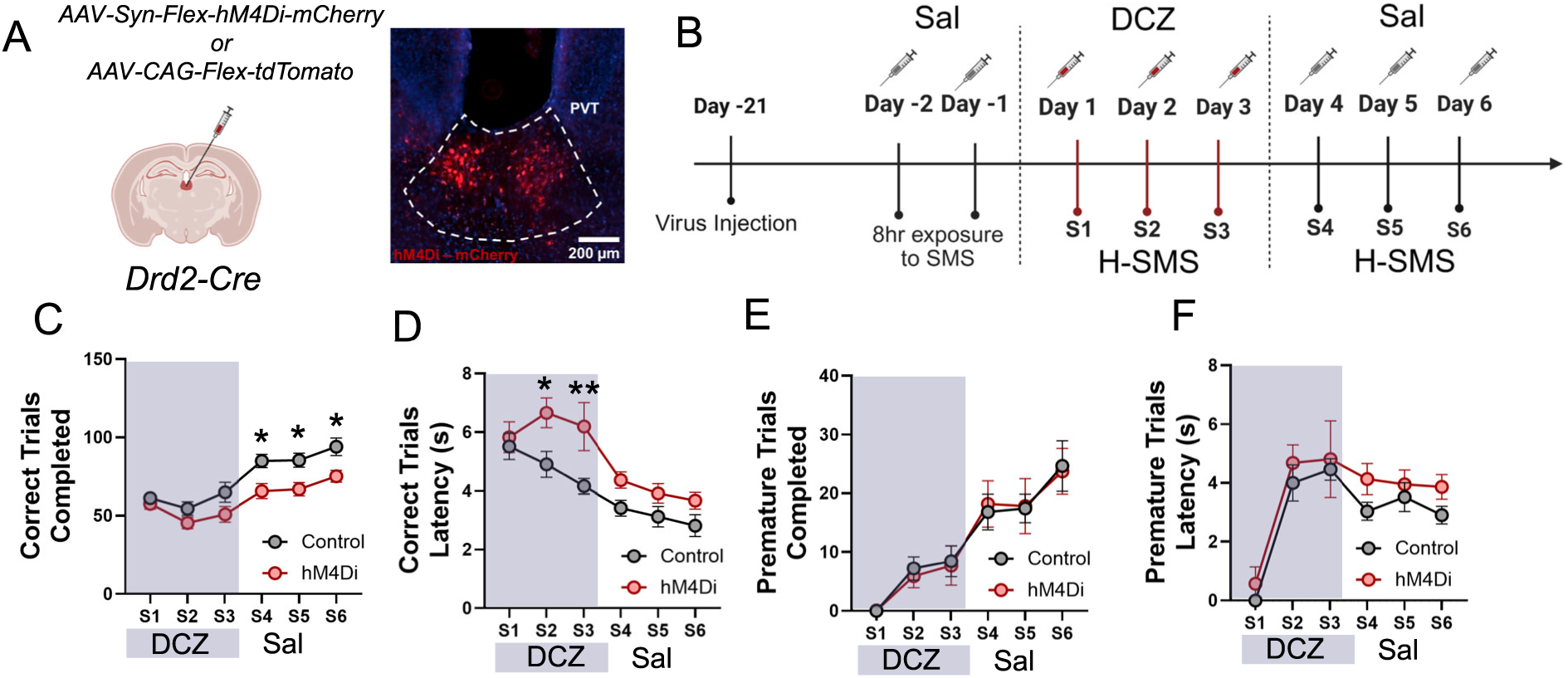
Chronic PVT^D2R+^ neuronal inhibition impairs performance in the linear maze task. **(A)** *Left*: Schematic of the stereotaxic injections for targeting D2R+ neurons in the PVT. *Right*: Representative image from a Drd2-Cre mouse expressing hM4Di-mcherry in the PVT. **(B)** Diagram showing the timeline for the chemogenetic inhibition of PVT^D2R+^ neurons. **(C)** Quantification of correct trials completed for hM4Di and Controls groups. Two-way ANOVA, Effect of test session \*\*\*\**p<0.*0001. Effect of viral group \*\*\*\**p<0.0001.* Interaction *p=0.47, ns*. Šídák’s multiple comparisons test: (S4) control vs. hM4Di, \**p<0.05;* (S5) control vs. hM4Di, \**p<0.05;* (S6) control vs. hM4Di, \**p<0.05.* **(D)** Quantification of latency to approach during correct trials for hM4Di and Controls groups. Two-way ANOVA, Effect of test session \*\*\*\**p<0.*0001. Effect of viral group \*\*\*\**p<0.0001;* Interaction *p=0.28, ns*. Šídák’s multiple comparisons test: (S2) control vs. hM4Di, \**p<0.05*; (S3) control vs. hM4Di, \*\**p<0.01.* **(E)** Quantification of premature trials completed for hM4Di and Controls groups. Two-way ANOVA, Effect of test session \*\*\*\**p<0.*0001. Effect of viral group \**p<0.05.* Interaction *p=0.99, ns*. **(F)** Quantification of latency to approach during premature trials for hM4Di and Controls groups. Two-way ANOVA, Effect of test session \*\*\*\**p<0.*0001. Effect of viral group \**p<0.05.* Interaction *p=0.29, ns*.

## DISCUSSION

In this study, we characterized a specific role for PVT^D2R+^ neurons in using interoceptive cues to guide goal-directed behavior. We showed that PVT^D2R+^ neurons compute an interoceptive prediction by integrating factors, such as current physiological needs, previous experiences, and environmental cues, to guide motivated behavior. Using a linear maze task that assesses motivated behavior in combination with *in vivo* fiber photometry of PVT^D2R+^ neurons, we found a significant increase in the response of these neurons during reward approach and reward delivery. Furthermore, these increases were directly related to the current state of the animal but also depend on the outcome expectation. We only observed increases in PVT^D2R+^ neuronal activity during motivated behavior of hungry animals expecting a food reward, but not during the motivated behavior of hungry animals expecting a water outcome, or during the return to the trigger zone. In addition, when mice were thirsty and a water outcome was provided, these neurons again showed increases during reward approach.

### Sensing of current physiological states by PVT neurons

Previous anatomical and functional studies have already established the PVT as a critical hub for sensing and responding to changes in homeostasis^15–17^. These studies highlighted the role of PVT neurons as key detectors of homeostatic imbalances, including changes in energy levels^18,29,44–49^, hydration^13,14,50^, stress^26–28,51^, pain^23–25^, thermoregulation^21,22^, itch^52,53^, social behavior^54–56^, and arousal states^57–59^. Anatomically, PVT neurons are richly connected to brain regions that monitor these homeostatic signals, such as the hypothalamus, which relays information about hunger, thirst, and stress through hormonal and neuropeptide signals, including orexin^18,60,61^, neuropeptide Y^19^, and others^62^. The PVT also shares rich connections with brainstem areas, such as rostral ventrolateral medulla^29^ and the nucleus of the solitary tract^47^, which receive information about the state of the body through somatic afferents and chemosensory receptors^11^.

Furthermore, PVT neurons express some of these chemosensory receptors, which enable them to directly sense and respond to homeostatic challenges. For example, PVT neurons express the Glut2 glucose transporter and are capable of directly detecting glucose levels^63^. Electrophysiological recordings show that glucose directly modulates the firing activity of PVT neurons, with some neurons exhibiting increased excitability in response to low glucose levels^63^. PVT neurons also sense changes in extracellular glucose concentrations via expression of enzymes like glucokinase^48^. Moreover, changes in hydration status, such as dehydration, activate PVT neurons, which in turn promote thirst-related behaviors^13,14^. Beyond basic survival needs, the PVT contributes to maintaining arousal and circadian rhythms, responding to dynamic changes in sleep-wake states and coordinating transitions between them^37,57,64^. Similarly, the PVT regulates stress responses via inputs from the hypothalamus and brainstem and projections to the amygdala and prefrontal cortex, influencing both emotional and physiological adaptations^26,27,33–35,65^. Experimental manipulation of PVT activity using optogenetics and chemogenetics further underscores its role in homeostasis. For instance, activating or inhibiting specific PVT subpopulations can alter feeding^18,31^, drinking^13,14^, stress reactivity^26,33,35^, and arousal levels^37,58^. Collectively, our findings are consistent with previous research showing that the magnitude of the activity of PVT neurons encodes the salience of current internal states, such as the degree of hunger or the level of stress. We also expand on this knowledge by showing that PVT^D2R+^ neurons are likely involved in these state-dependent processes.

Notably, in our study, we also found that a non-caloric reward showed increased PVT^D2R+^ neuronal responses during reward approach and reward delivery for both hunger and sated states. This was somewhat surprising given that a non-caloric reward presumably does not relay information about nutritional value or physiological significance. However, when these non-caloric rewards were provided, the within-session state-dependent modulation was no longer observed, and this effect persisted even with prolonged exposure to the non-caloric solution. Other studies have also found differences in neurocircuit involvement following non-caloric sweetener consumption. The intake of the non-caloric sweetener, sucralose, fails to deactivate hypothalamic areas compared to the ingestion of glucose and fructose^66^. Additionally, compared to caloric options, sucralose consumption shows reduced activity in areas related to reward, including the lateral hypothalamus (LHA), ventral tegmental area (VTA), and nucleus accumbens (NAc)^67^. However, despite their wide use, relatively little is known about the effects of non-caloric sweeteners in the brain. Indeed, one of the suggested potential risks of non-caloric food intake is the decoupling of sweet taste from the satiety effects needed for maintaining the finely tuned regulation of energy homeostasis^68^. Future research should investigate the underlying mechanisms of how sweet tastants activate PVT^D2R+^ neurons and the observed decoupling of reward and satiety we observed.

### Modulation of PVT^D2R+^ neurons by environmental cues and context

Previous studies established that PVT neurons are particularly sensitive to external signals that predict outcomes critical for survival, such as food, safety, or danger^18,19,27,32,33,35,47,59^. When an environmental cue is associated with a physiologically relevant outcome, PVT neurons become activated by the predictive cues^18,33^. This activation is proposed to facilitate the integration of relevant sensory information, to enable the prediction of future internal physiological states to drive adaptive responses accordingly. For instance, in a hunger state, the PVT exhibits heightened activity in response to food-related cues^18^. This heightened response to physiologically relevant cues has been documented in other states like thirst^13,14^, fear^26,33,35^, safety^27^, and social^55^ behaviors. These responses engage with downstream brain regions like the NAc^18,26,29,39^, amygdala^24,33,35^ and prefrontal cortex^27,30,33,34^ to prioritize behaviors that will reestablish homeostasis^13,15,29,69^. We confirmed and expanded this previous knowledge by showing that cues and context modulated the responses of PVT^D2R+^ neurons, as shown by the differential responses to the reward zone approach vs. trigger zone return (Fig. 1 and Fig. 6) and correct vs. premature trials (Fig. 4O). These context and cue-driven activations reflect the ability of PVT^D2R+^ neurons to dynamically adjust their activity based on the motivational significance of the stimulus, ensuring that resources are directed toward the most urgent physiological needs. Additionally, a major finding of this study is the slow emergence of these cue-induced responses, which are in contrast to the fast emergence of cue-induced responses in other brain regions^33,34,70^. Indeed, the slow emergence of cue related responses in PVT neurons has been previously documented in the literature^33,34^ and are in accordance with our new proposed role of PVT^D2R+^ neurons in driving interoceptive predictions. This is consistent with modern interoceptive models, which posit that predictive interoceptive responses to environmental cues tend to emerge at a much slower pace than exteroceptive predictions^4,5,9^. This slow emergence has been attributed to interoceptive signals being more subtle and diffused than exteroceptive ones^4,5^. Importantly, dysregulation of cue-driven activation has been implicated in conditions like substance use disorder, where maladaptive cue-responses lead to compulsive behaviors. This highlights the critical balance the PVT^D2R+^ neurons might maintain in the processing and prioritizing of environmental stimuli.

### Modulation of PVT^D2R+^ neurons by learned experiences and their role in interoceptive predictions

Models of interoception also posit that beyond sensing of current internal signals from the body, the brain also makes predictions of future internal states^3–6,9^. Indeed, these same models postulate that predictive coding of internal states are not represented in a single brain area but in the connectivity of various hierarchical populations of neurons. This hierarchy enables representation at various levels of abstraction, with higher levels integrating information across multiple sensory domains to create more abstract predictions, while lower levels integrate more exact sensory inputs^9^. Previous studies in humans and rodents show that one key higher-level area involved in these predictions is the insula cortex^1,50,71^. Importantly, the PVT shares strong anatomical connections with the insula cortex that participate in the predictive coding of future internal states^50,72^. Here, we also expand on this knowledge to show that PVT^D2R+^ neurons are part of the hierarchy of populations making predictions of internal states. We now show that beyond their role in detecting current internal states, PVT^D2R+^ neurons also dynamically adjust their activity to reflect both the internal state of the organism and the expected value of the outcome. This was most obviously demonstrated by the increased PVT^D2R+^ neuronal activity observed when mice were approaching the reward zone. Conversely, the same motivated behavior, ‘running’ towards the trigger zone, resulted in decreases in the responses of these neurons. Furthermore, these reductions in photometric responses of PVT^D2R+^ neurons during the return phase were also associated with enhanced GABAergic inhibitory signaling. Indeed, the PVT receives diverse and wide-ranging direct inhibitory inputs from brain areas such as the zona incerta, anterior hypothalamic nucleus, and periaqueductal gray, among others^26^. Additionally, it receives indirect inhibitory projections from the prelimbic cortex through the anterior ventral thalamic reticular nucleus^73,74^. Recent previous studies showed that this indirect pathway is critical for shaping goal-directed decisions through a disinhibitory mechanism^73,74^. Importantly, and consistent with their ability to make interoceptive predictions, the responses during the decision to return were influenced by the state of the animal and the expected outcome. In addition, analyzing premature approaches allowed us to assess the responses of PVT^D2R+^ neurons while animals ran with the same vigor and in the same context (toward the reward zone) but without a cue light signaling the reward. During these premature approaches, the activity of PVT^D2R+^ neurons was attenuated. Moreover, these differences between correct and premature responses were not observed when the outcome was no longer relevant (Hungry-H_2_O sessions and Hungry-OM sessions). These findings strongly suggest that when the outcome is not relevant, PVT^D2R+^ neurons adjust their activity to update future interoceptive predictions.

### Interoception deficits as a major underlying mechanism for prominent psychiatric conditions

Interoception is one of the core drivers of motivation and emotion^2–4,6,8^. However, when interoceptive systems become dysregulated, it can lead to motivational and emotional deficits seen in many prominent psychiatric conditions, including chronic stress, major depressive disorder (MDD), autism spectrum disorders (ASDs), and others^2,3,6,8^. Both human and rodent studies show that the paraventricular thalamus (PVT) contributes to these disorders in different ways. For example, chronic stress manifests physiologically with increases in cortisol levels and/or impaired hypothalamus-pituitary-adrenal (HPA) axis activation^75,76^. PVT activity is altered after chronic stress^26,77^, and lesions or inactivation of the PVT impair animals’ ability to adapt to novel stressors^28^. MDD, characterized by low mood and low motivation, is highly linked to autonomic dysfunction and inflammation^78^. Studies in animals show that inflammation results in PVT dysfunction and motivational impairments^79–82^, indicating that dysregulation of PVT neurons could be a key driver of low motivation in MDD. Additionally, anxiety disorders, eating disorders, substance use disorders, and pain disorders are other well-known conditions resulting from dysregulation of interoceptive processes^2,3,6,8^. The PVT has been implicated in these conditions in both humans and rodents^83^.

### Limitations

Previous studies have demonstrated that PVT neurons exhibit heterogeneity, which has important functional implications^37–39^. For example, our previous research has shown that other types of neurons, such as PVT neurons that do not express the D2 receptor, are functionally distinct from PVT^D2R+^ neurons. During motivated behavior, these PVT^D2R-^ neurons show decreases in photometric responses and are not as attuned to the physiological state of the animal^39^. Therefore, one main limitation of the present study is the use of fiber photometry to investigate the activity of PVT^D2R+^ neurons and their role in interoceptive signaling and predictions. It is possible that the observed PVT^D2R+^ neuronal activity shown here includes responses from other PVT subpopulations within the PVT^D2R+^ neurons. Indeed, using single nucleus RNA, a previous study showed that PVT^D2R+^ neurons can be further subdivided into two subtypes: Esr1+ and Col12a1+^38^. Thus, it is possible that the distinct effects observed are mediated by these distinct subpopulations of PVT^D2R+^ neurons. Due to these limitations, future studies should use single-cell calcium imaging of PVT neurons to fully characterize the activity of PVT^D2R+^ and PVT^D2R-^ neurons during interoceptive prediction. This approach will help understand how factors such as physiological states, environmental cues, and previous experiences impact these neurons and their role in interoceptive predictions.

In conclusion, pinpointing the functional role of PVT neurons and their contribution to motivated behavior has been challenging. This difficulty arises from the recently discovered heterogeneity in PVT neurons^37–39,84^ and their opposing functional responses^39^. Additionally, the PVT has been implicated in various conditions and roles, adding to the complexity. The findings presented here illuminate the role of PVT^D2R+^ neurons in mediating interoceptive predictions, proposing a more unified role that contextualizes previous findings. Collectively, and as highlighted here, interoception signaling and predictions are central to creating internal states that guide adaptive behavior. Dysregulation of interoceptive processes results in a range of psychiatric and neurological conditions. Thus, a deeper understanding of the neural basis of interoception may offer new insights into potential therapeutics for these debilitating conditions. By elucidating key areas and neuronal populations involved and how they respond to different factors, this research may help identify new therapeutic targets aimed at restoring healthy interoceptive signaling in individuals.

## ACKNOWLEDGEMENTS

We thank Dr. Zachary Knight, Dr. Kirstie Cummings, Dr. Jamie Peters, Dr. Barry Setlow, Dr. Chase Francis and Jonathan Ross Folmar for their comments on the manuscript. This work was supported by Department of Neurobiology at the University of Alabama Birmingham, an R00 MH126429 to S.B., a NARSAD Young Investigator Award by the Brain and Behavior Research Foundation to S.B., a T32 by the National Cancer Institute of the National Institutes of Health under award number T32CA047888 to B.M., a NIGMS K12 Institutional Research and Career Development Award to S.N.M. and by the NIMH Intramural Research Program (project ZIC-MH002968, authors A.X., F.P., and G.L.).

## AUTHOR CONTRIBUTIONS

B.M., C.L., and S.N.M. performed the behavioral experiments and surgeries. L.A., J.T., and S.H. assisted with behavioral experiments and histology. B.M. and S.B. performed all the data analysis. A.X., G.L., and F.P. assisted with the FLMM concurrent model analyses. S.B. designed the study and interpreted the results. S.B. wrote the manuscript. B.M., S.N.M., and G.L., provided extensive edits and comments to the manuscript.

---

**Supplementary Figure 1. PVTD2R^+^ neurons sense and track hunger states. (A)** Schematic depicting the linear maze reward-seeking task. **(B)** *Left:* Average GCaMP6s responses from PVT^D2R+^ neurons showing ramping activity during reward approach. *Right:* Quantification of the approach-evoked changes in GCaMP6s fluorescence in PVT^D2R+^ neurons. Area under the curve (AUC), n = 705 trials from 5 mice. Comparisons between baseline (base) and approach (H-SMS); two-tailed paired t-test, *\*\*\*\*p<0.0001*. **(C)** Heatmap showing excitatory reward approach responses from PVT^D2R+^ neurons, time-locked to approach onset, sorted by trial order, and binned into 5 ‘trial group blocks’ (G1 – G5). **(D)** Latencies to reach the reward zone across ‘trial group blocks’ (G1 – G5). Repeated measures ANOVA, *\*\*p<0.01*; G1 vs. G5 Dunnett’s multiple comparisons test *p=0.98*, *ns.* **(E)** Average GCaMP6s responses from PVT^D2R+^ neurons comparing reward approach between early ‘G1’ and late ‘G5’ trials within the test sessions. **(F)** AUC quantification of the reward approach-evoked changes in GCaMP6s activity across trial group blocks. Repeated measures ANOVA, *\*\*\*p<0.001*. G1 vs. G5 Dunnett’s multiple comparisons test *\*p<0.05*. **(G)** Correlation between the AUC of the reward approach-evoked changes in GCaMP6s activity and trial order within a test session. **(H)** FLMM coefficient estimates plot of the approach trial order effect and statistical significance at each trial time-point for the photometric responses of PVT^D2R+^ neurons for reward zone entry. The plot shows a negative association between PVT^D2R+^ GCaMP6s responses and trial order before and after reward zone entry. **(I)** No correlation between the AUC of the reward approach-evoked changes in GCaMP6s activity and approach latency within a test session.

**Supplementary Figure 2. Long-term exposure to the non-caloric reward did not affect the magnitude of PVT^D2R+^ neurons’ photometric responses.**

**(A)** Timeline schematic for non-caloric reward exposure and test sessions. **(B)** Average GCaMP6s responses (n=5) from PVT^D2R+^ neurons for the non-caloric reward after two weeks of exposure. **(C)** Average GCaMP6s responses from PVT^D2R+^ neurons comparing reward approach between early ‘G1’ and late ‘G5’ trials within the test sessions. **(D)** AUC quantification of the reward approach-evoked changes in GCaMP6s activity across trial group blocks. Repeated measures ANOVA, *p=0.39*, *ns*. G1 vs. G5 Dunnett’s multiple comparisons test *p=0.50, ns*.

**Supplementary Figure 3. Premature trials are associated with reduced photometric responses of PVT^D2R+^ neurons during reward approach.**

**(A)** Average GCaMP6s responses (n=5) from PVT^D2R+^ neurons and AUC quantification showing comparisons between ramping activity during reward approach in correct and premature trials in the H-SMS test sessions. Two-tailed paired t-test \*\**p<0.01*. **(B)** Latencies to reach the reward zone for correct and premature trials performed in the H-SMS test sessions. Two-tailed paired t-test *p=0.60, ns*. **(C)** AUC quantification of the reward approach-evoked changes in GCaMP6s activity across trial group blocks for the H-SMS premature trials. Repeated measures ANOVA, \*\*\**p<0.001*. G1 vs. G5 Dunnett’s multiple comparisons test \*\**p<0.01.* **(D)** AUC quantification of the reward approach across ‘trial group blocks’ (G1 – G5) for H-SMS-C (correct) and H-SMS-P (premature) trials. Two-way ANOVA, Effect of trial group block *\*\*\*\*p<0.0001*. Effect of trial type *\*\*\*\*p<0.0001;* Interaction *p=0.61, ns*.

**Supplementary Figure 4. Concurrent FLMM model of between conditions comparisons.**

**(A-E)** FLMM coefficient estimates plots and statistical significance at each trial time-point for the mean changes in photometric responses of PVT^D2R+^ neurons during approach and reward delivery for the between outcomes comparisons (i.e., H-SMS vs. ‘other outcome’). Each plot shows the four beta coefficients that result from these comparisons. *Top left* - β*_0_(s):* shows the mean signal when the mouse received the ‘other outcome’. *Top right* - β*_1_(s):* shows the mean difference in signal between H-SMS and the ‘other outcome’ during reward delivery. *Bottom left* - β*_2_(s):* shows the mean difference between approach and outcome delivery for the ‘other outcome’. *Bottom right* - β*_3_(s):* shows the difference of differences, capturing the additional effect on signal when the mouse is approaching the ‘other outcome’ that is not already captured by β*_0_(s)+*β*_1_(s)+*β*_2_(s).* As such, β*_3_(s)* captures the interaction effect between behavioral state (approaching vs. reward delivery) and outcome condition (H-SMS vs. ‘other outcome’).

**(A)** Beta coefficients for the comparisons of the H-SMS and S-SMS conditions. β*_0_(s):* shows that during reward delivery in the S-SMS condition, the mean signal is not significantly modulated. β*_1_(s):* shows that during reward delivery, the mean signal is significantly higher for H-SMS compared to S-SMS conditions. β*_2_(s):* shows that the mean signal during approach is significantly lower than the mean signal during reward delivery for the S-SMS condition. β*_3_(s):* shows a significant negative interaction effect between behavioral states (approaching vs. receiving) and conditions (H-SMS vs. S-SMS), showing that the mean changes in photometric responses are larger during the outcome delivery than during approach. Specifically, there are larger mean changes during H-SMS than in the S-SMS conditions.

**(B)** Beta coefficients for the comparisons of the H-SMS and H-NCal conditions. β*_0_(s):* shows that during reward delivery in the H-NCal condition, the mean signal is significantly positively modulated by the NCal rewards. β*_1_(s):* shows that during reward delivery, the mean signal is significantly lower for H-SMS compared to H-NCal conditions. β*_2_(s):* shows that the mean signal during approach is significantly lower than the signal during reward delivery for the H-NCal condition. β*_3_(s):* shows a significant negative interaction early in the trial, followed by positive interaction effects later in the trial and during reward delivery. This indicates that the differences in neural activity during the early approach are significantly lower, but later in the trial, and during reward consumption, they are significantly higher in the NCal than in the H-SMS conditions.

**(C)** Beta coefficients for the comparisons of the H-SMS and H-Suc conditions. β*_0_(s)*: shows that during reward delivery in the H-Suc condition, the mean signal is positively modulated by the Suc rewards, however, this was not significant. β*_1_(s):* shows that during reward delivery, the mean signal for H-SMS is not significantly different than the signal during the H-Suc conditions. β*_2_(s):* shows that the mean signal during approach is significantly lower than the signal during reward delivery for the H-Suc condition. β*_3_(s):* shows no significant interaction effects between behavioral states (approaching vs. receiving) and conditions (H-SMS vs. H-Suc).

**(D)** Beta coefficients for the comparisons of the H-SMS and H-H_2_O conditions. β*_0_(s):* shows that during reward delivery in the H-H_2_O condition, the mean signal is not significantly modulated. β*_1_(s):* shows that during reward delivery, the mean signal is significantly higher for H-SMS compared to H-H_2_O conditions. β*_2_(s):* shows that the mean signal during approach is significantly lower than the signal during outcome delivery for the H-H_2_O condition. β*_3_(s):* shows a significant negative interaction effect between behavioral states (approaching vs. receiving) and conditions (H-SMS vs. H-H_2_O), showing that the mean changes in photometric responses are larger during the outcome delivery than during approach. Specifically, there are larger mean changes during H-SMS than in the H-H_2_O conditions.

**(E)** Beta coefficients for the comparisons of the H-SMS and H-OM conditions. β*_0_(s):* shows that during reward delivery in the H-OM condition, the mean signal is positively modulated initially by the omission of the rewards, but the signal quickly goes down over time. β*_1_(s):* shows that during reward delivery, the mean signal is significantly higher for H-SMS compared to H-OM conditions. β*_2_(s):* shows that the mean signal during approach is significantly lower than the signal during outcome delivery for the H-OM condition. β*_3_(s):* shows a significant negative interaction effects in the trial and during reward delivery.

**Supplementary Figure 5. Ti**me **spent in the reward zone after reward delivery does not differ across outcome groups or trial groups.**

**(A-F)** Quantifications of time spent in the reward zone after reward delivery and prior to return across trial groups for the **(A)** H-SMS test sessions; Repeated measures ANOVA, *p=0.07, ns.* G1 vs. G5 Dunnett’s multiple comparisons test *p=0.52, ns*. **(B)** S-SMS test sessions; Repeated measures ANOVA, *p=0.11, ns.* G1 vs. G5 Dunnett’s multiple comparisons test *p=0.07, ns*. **(C)** H-NCal test sessions; Repeated measures ANOVA, *p=0.16, ns.* G1 vs. G5 Dunnett’s multiple comparisons test *p=0.77, ns*. **(D)** H-Suc test sessions; Repeated measures ANOVA, *p=0.44, ns.* G1 vs. G5 Dunnett’s multiple comparisons test *p=0.95, ns*. **(E)** H-H_2_O test sessions; Repeated measures ANOVA, *p=0.11, ns.* G1 vs. G5 Dunnett’s multiple comparisons test *p=0.06, ns*. **(F)** H-OM test sessions; Repeated measures ANOVA, *p=0.51, ns.* G1 vs. G5 Dunnett’s multiple comparisons test *p=0.99, ns*. **(G)** Quantifications of time spent in the reward zone after reward delivery and prior to return across outcome groups. Repeated measures ANOVA, *p=0.26, ns.* H-SMS vs. S-SMS Dunnett’s multiple comparisons test *p=0.99, ns*. H-SMS vs. H-NCal Dunnett’s multiple comparisons test *p=0.63, ns*. H-SMS vs. H-Suc Dunnett’s multiple comparisons test *p=0.99, ns*. H-SMS vs. H-H_2_O Dunnett’s multiple comparisons test *p=0.32, ns*. H-SMS vs. H-OM Dunnett’s multiple comparisons test *p=0.49, ns*.

**Supplementary Figure 6. GABAergic signaling dynamics in PVT neurons during performance in the linear maze task.**

**(A)** Schematic of the stereotaxic injections and optical fiber implantation for targeting PVT and PVT^D2R+^ neurons. **(B)** Representative image from a C57BL/6 mouse expressing the GABA sensor iGABASnFR2 in the PVT. **(C)** Timeline schematic for the test sessions. **(D-G)** Average iGABASnFR2 responses from PVT neurons and PVT^D2R+^ neurons during reward approach and during return in H-SMS test sessions. PVT neuronal imaging: 1474 trials from 5 mice. PVT^D2R+^ neuronal imaging: 1283 trials from 5 mice. **(H-K)** Average iGABASnFR2 responses from PVT and PVT^D2R+^ neurons comparing reward approach and return between early ‘G1’ and late ‘G5’ trials within the H-SMS test sessions. **(L-O)** AUC quantification of the reward approach-evoked changes and return-evoked changes in iGABASnFR2 responses from PVT neurons and PVT^D2R+^ neurons activity across trial group blocks. **(L)** Repeated measures ANOVA, *p=0.30*. G1 vs. G5 Dunnett’s multiple comparisons test *p=0.22, ns*. **(M)** Repeated measures ANOVA, *p=0.66*. G1 vs. G5 Dunnett’s multiple comparisons test *p=0.63, ns*. **(N)** Repeated measures ANOVA, *p=0.22, ns*. G1 vs. G5 Dunnett’s multiple comparisons test *p=0.23, ns*. **(O)** Repeated measures ANOVA, *p=0.18, ns*. G1 vs. G5 Dunnett’s multiple comparisons test *p=0.57, ns*.

**Supplementary Figure 7. PVTD2R^+^ photometric responses increased during the decision to return, and these increases were modulated by state and outcome expectation.**

**(A)** Representative single trace of the photometric responses of PVT^D2R+^ neurons at the different stages of the trial. **(B)** Quantification of the maximum peak responses during decision to return (1 second before return)-evoked PVT^D2R+^ GCaMP6s activity for sessions H-SMS (strong responses modulated by state), S-SMS (weak responses modulated by state), H-NCal (strong responses not modulated by state), and H-OM (weak responses not modulated by state). Repeated measures ANOVA, \*\**p<0.01*. H-SMS vs. S-SMS Dunnett’s multiple comparisons test *p=0.31, ns*. H-SMS vs. H-NCal Dunnett’s multiple comparisons test *p=0.99, ns*. H-SMS vs. H-OM Dunnett’s multiple comparisons test \*\**p<0.01*. **(C-F)** Quantification of the maximum peak responses during decision to return-evoked PVT^D2R+^ GCaMP6s activity across trial group block for the H-SMS, S-SMS, H-NCal, and H-OM test sessions. **(C)** H-SMS: Repeated measures ANOVA, \*\*\**p<0.001*. G1 vs. G5 Dunnett’s multiple comparisons test \**p<0.05*. **(D)** S-SMS: Repeated measures ANOVA, \*\*\**p<0.001*. G1 vs. G5 Dunnett’s multiple comparisons test \*\**p<0.01*. **(E)** H-NCal: Repeated measures ANOVA, *p=0.11, ns*. G1 vs. G5 Dunnett’s multiple comparisons test *p=0.86, ns*. **(F)** H-OM: Repeated measures ANOVA, *p=0.69, ns*. G1 vs. G5 Dunnett’s multiple comparisons test *p=0.99, ns*. **(G-J)** Correlations between the maximum peak in signal 1 sec before return and the trial order for H-SMS, S-SMS, H-NCal, and H-OM test sessions.

**Supplementary Figure 8. Motivational states are associated with PVT^D2R+^ neuronal responses across different conditions.**

**(A-G)** Average GCaMP6s responses from PVT^D2R+^ neurons comparing between fast ‘L1’ and slow ‘L5’ trials (top). AUC quantifications across latency groups during approach (middle). Correlations between the approach slope and the latency to approach (bottom) for the **(A)** H-SMS test sessions; Repeated measures ANOVA, \*\**p<0.01.* L1 vs. L5 Dunnett’s multiple comparisons test \*\**p<0.01*. **(B)** S-SMS test sessions; Repeated measures ANOVA, *p=0.06, ns.* L1 vs. L5 Dunnett’s multiple comparisons test \*\**p<0.01*. **(C)** H-NCal test sessions; Repeated measures ANOVA, \**p<0.05.* L1 vs. L5 Dunnett’s multiple comparisons test \**p<0.05*. **(D)** H-Suc test sessions; Repeated measures ANOVA, \*\**p<0.01.* L1 vs. L5 Dunnett’s multiple comparisons test \**p<0.05*. **(E)** H-H_2_O test sessions; Repeated measures ANOVA, *p=0.62, ns.* L1 vs. L5 Dunnett’s multiple comparisons test *p=0.95, ns*. **(F)** H-OM test sessions; Repeated measures ANOVA, \*\**p<0.01.* L1 vs. L5 Dunnett’s multiple comparisons test \*\*\**p<0.001*. **(G)** S-OM test sessions; Repeated measures ANOVA, *p=0.57, ns.* L1 vs. L5 Dunnett’s multiple comparisons test *p=0.99, ns*.

**Supplementary Figure 9. PVTD2R^+^ responses in individual animals during approach and return over learning.**

**(A)** Mean GCaMP6s responses of PVT^D2R+^ neurons during approach, per animal, across learning. **(B)** Mean GCaMP6s responses of PVT^D2R+^ neurons during return, per animal, across learning.

**Supplementary Figure 10. Chronic, but not acute, chemogenetic inhibition of PVT^D2R+^ neurons via CNO administration results in impaired performance on the linear maze task.**

**(A)** *Left:* Schematic of the stereotaxic injections for targeting D2R+ neurons in the PVT. *Right:* Representative image from a Drd2-Cre mouse expressing hM4Di-mcherry in the PVT. **(B)** Schematic of the timeline for the chemogenetic chronic inhibition of PVT^D2R+^ neurons with CNO.

**(C)** Quantification of correct trials completed for hM4Di and Controls groups. Two-way ANOVA, Effect of test session *\*\*\*\*p<0.0001*. Effect of viral group *\*\*p<0.01*. Interaction \**\*\*p<0.001*. *Šídák’s multiple comparisons test: (S5) control vs. hM4Di, *p<0.05; (S6) control vs. hM4Di, ***p<0.001.* **(D)** Quantification of the latency to approach during correct trials for hM4Di and Controls groups. Two-way ANOVA, Effect of test session *\*\*\*\*p<0.0001*. Effect of viral group *\*\*\*p<0.001*; Interaction *p=0.50, ns.* Šídák’s multiple comparisons test: (S3) control vs. hM4Di, *\*p<0.05*. **(E)** Quantification of premature trials completed for hM4Di and Controls groups. Two-way ANOVA, Effect of test session *\*\*\*\*p<0.0001*. Effect of viral group *p=0.91,ns*. Interaction *p=0.86, ns*. **(F)** Quantification of latency to approach during premature trials for hM4Di and Controls groups. Two-way ANOVA, Effect of test session *\*p<0.05*. Effect of viral group *\*\*p<0.01*. Interaction *p=0.24*, ns. Šídák’s multiple comparisons test: (S2) control vs. hM4Di, *\*\*p<0.01*. **(G)** Schematic of the stereotaxic injections for targeting D2R+ neurons in the PVT. **(H)** Schematic of the timeline for the chemogenetic acute inhibition of PVT^D2R+^ neurons with CNO**. (I)** Quantification of correct trials completed for hM4Di and Controls groups. Two-way ANOVA, Effect of test session *p=0.07, ns*. Effect of viral group *p=0.73, ns*. Interaction *p=0.88, ns*. **(J)** Quantification of premature trials completed for hM4Di and Controls groups. Two-way ANOVA, Effect of test session *p=0.05, ns.* Effect of viral group *p=0.34, ns.* Interaction *p=0.98, ns*.

